# Mutational scanning of TnpB reveals latent activity for genome editing

**DOI:** 10.1101/2025.02.11.637750

**Authors:** Brittney W. Thornton, Rachel F. Weissman, Jorge E. Rodriguez, Ryan V. Tran, Brenda T. Duong, Cynthia I. Terrace, Ugrappa Nagalakshmi, George Austin, Evan D. Groover, Flora Zhiqi Wang, Jung-Un Park, Viktoriya Georgieva, Julia Tartaglia, Myeong-Je Cho, Savithramma P. Dinesh-Kumar, Jennifer A. Doudna, David F. Savage

## Abstract

TnpB is a diverse family of RNA-guided endonucleases associated with prokaryotic transposons. Due to their small size and putative evolutionary relationship to CRISPR-Cas12, TnpB enzymes hold significant potential for genome editing. However, most TnpBs lack robust gene editing activity, and unbiased profiling of mutational effects on editing activity has not been explored. Here, we mapped comprehensive sequence-function landscapes of a TnpB ribonucleoprotein and discovered many activating mutations in both the protein and RNA. One- and two-position RNA mutants outperform existing variants, highlighting the utility of systematic RNA scaffold mutagenesis. Leveraging the protein’s mutational landscape, we identified enhanced TnpB variants from a combinatorial library of activating mutations. These variants enhanced editing in human cells, *N. benthamiana*, pepper, and rice, with up to a fifty-fold increase compared to wild-type TnpB. These findings highlight previously unknown elements critical for regulating TnpB endonuclease activity and reveal surprising latent activity accessible through mutation.

## Introduction

TnpBs are a family of RNA-guided endonuclease proteins encoded within IS200/IS605 and IS607 transposons and are thought to be the evolutionary ancestors of Type V CRISPR-Cas enzymes^1–5^. TnpB binds to the right end element RNA (reRNA), which has a noncoding 5’ scaffold and a 3’ variable region, to guide ribonucleoprotein (RNP) cleavage at a complementary DNA sequence proximal to a transposon-associated motif (TAM)^2,3^. As a putative evolutionary predecessor to the CRISPR-Cas12 enzymes, TnpB retains core domains shared across this CRISPR protein family^4–7^. Understanding the relationship between TnpB’s protein sequence and activity can provide both fundamental knowledge and serve as a basis for engineering improved or altered RNA-guided endonucleases.

While highly active variants of CRISPR-Cas enzymes have been identified through protein engineering and rational design^8^, these approaches often explore a limited sequence space. The conformational changes TnpB undergoes during the dynamic coordination of nucleic acid binding, catalytic center activation, and DNA cleavage^6,7^ make it challenging to predict the effects of mutations. Deep mutational scanning (DMS) approaches typically assess every individual amino acid mutation using high-throughput assays of protein function^9–11^. While DMS effectively provides comprehensive maps of protein function, it is often practically limited by protein size. TnpB is uniquely well-suited for this approach due to its compact amino acid length and RNA scaffold.

We conducted DMS over the entire ISDra2 TnpB RNP, one of the first experimentally characterized TnpB orthologs^2^. Utilizing a positive selection assay for DNA cleavage, we identified a broad spectrum of enhancing, neutral, and deleterious mutations within the TnpB protein and its reRNA. These data elucidate dynamic regions involved in DNA binding and cleavage, including a mutational hot-spot within the reRNA secondary structure where mutations increase DNA cleavage activity. We found that 20% of single amino acid mutations, many of which are not frequently observed in nature, increase activity relative to the wild-type (WT) TnpB protein. This suggests that native ISDra2 TnpB activity may be subject to negative selection, possibly due to its role as a transposon-associated homing endonuclease^2,12^. Furthermore, we identified combinations of activating mutations that increase TnpB-mediated genome editing activity in both human cells and plants.

## Results and Discussion

### Selection for TnpB-mediated DNA cleavage in yeast

In their native genomic context, the TnpB reRNA and protein-coding sequences overlap with each other and with insertion sequence elements essential for transposition, imposing unknown sequence-function constraints^2,3^. To interrogate the effects of mutations in both reRNA and TnpB protein without the native sequence constraints of transposition, we encoded the reRNA and codon-optimized ISDra2 TnpB protein under the control of separate regulatory elements (Fig. 1a). To assess on-target cleavage activity we adapted an *in vivo* selection previously used to enhance CRISPR-Cas9 activity, which utilizes yeast (*S. cerevisiae*) strains with a genomic *ade2*^-^ reporter cassette^13^ (Fig. 1b). On-target cleavage of the reporter cassette initiates *ADE2* repair, enabling cells to grow on media lacking adenine. Thus, reporter strain growth in the presence and absence of adenine can be used as a readout for target site cleavage (Fig. 1c). We first demonstrated that this assay can quantitatively measure endonuclease activity beyond that of CRISPR-Cas9, including WT ISDra2 TnpB and CRISPR-Cas12 endonucleases^13^ (Extended Data Fig. 1a).

**Figure 1.**
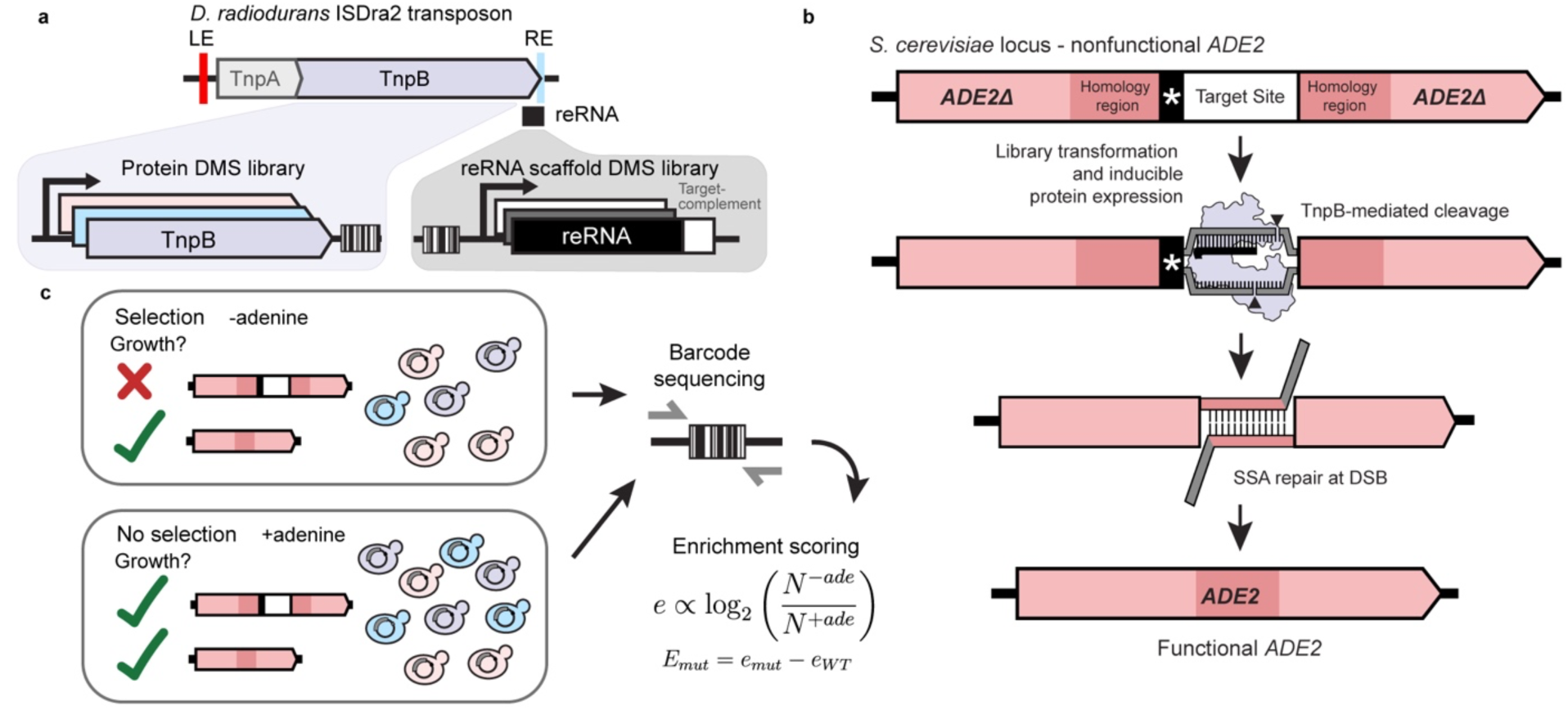
Design of deep mutational scanning libraries and optimized *in vivo* selection for endonuclease activity in yeast. **a,** TnpB protein and reRNA from the *D. radiodurans* ISDra2 transposon were placed under the control of separate regulatory elements. Deep mutational scanning libraries were constructed for both molecules and assayed separately. **b,** Schematic of the yeast-based cleavage assay used for individual variant testing and high-throughput library experiments. On-target double-stranded breaks in the reporter cassette enable repair of the *ADE2* locus by single-stranded annealing at duplicate homology regions. *ADE2* repair rescues colony growth in selective, adenine-deficient media. **c,** Representation of the library selection. Plasmid DNA from yeast grown in selective (-adenine) and non-selective (+adenine) conditions was extracted. Barcodes from plasmids were sequenced and the log-ratio of barcode abundance in selective over nonselective conditions was used to assess variant enrichment. Enrichment of variants was normalized to the wild-type TnpB RNP enrichment.

We constructed independent pooled plasmid libraries of reRNA and protein variants (Fig. 1a). The reRNA and protein DMS libraries were barcoded such that each variant was associated with ∼30 unique barcodes, providing statistical replicates. The libraries were transformed into yeast reporter strains with either their WT reRNA or protein counterparts, and barcode abundance in selective and nonselective media was quantified at multiple timepoints across two biological replicates. Relative variant enrichment was calculated as the log ratio of variant abundance in selective and non-selective conditions, and all enrichments were normalized to wild-type controls (Fig. 1c).

### Profiling the mutational landscape of the TnpB reRNA

The reRNA accounts for nearly half of the molecular weight of the ISDra2 TnpB RNP complex^6,7^. Previously, a truncation within the reRNA’s stem 2 region, termed Trim2, has been shown to maintain or increase ISDra2 TnpB’s genome editing activity^7,14^ (Extended Data Fig. 1b). Given the complex secondary and tertiary structures of the TnpB RNA scaffold, we hypothesized that comprehensive mutagenesis of the reRNA scaffold could reveal insights into evolutionary constraints on the reRNA sequence and RNP endonuclease activity.

The 116 nt reRNA scaffold is necessary and sufficient for TnpB-mediated cleavage^7^ and was chosen as the starting sequence (termed WT reRNA) for deep mutational scanning (Extended Data Fig. 1c). To investigate the mutational tolerance of the reRNA, our DMS library included all single-nucleotide substitutions, as well as both single- and double-nucleotide deletions. We also included variants with the disordered regions of stem 1 and stem 2 replaced with thermodynamically stable tetraloops^15,16^, the reported Trim2 variant, and variants with reported inactivating truncations within the triplex and pseudoknot serving as negative controls^14^, for a total of 576 assayed mutants (Fig. 2a-b).

**Figure 2.**
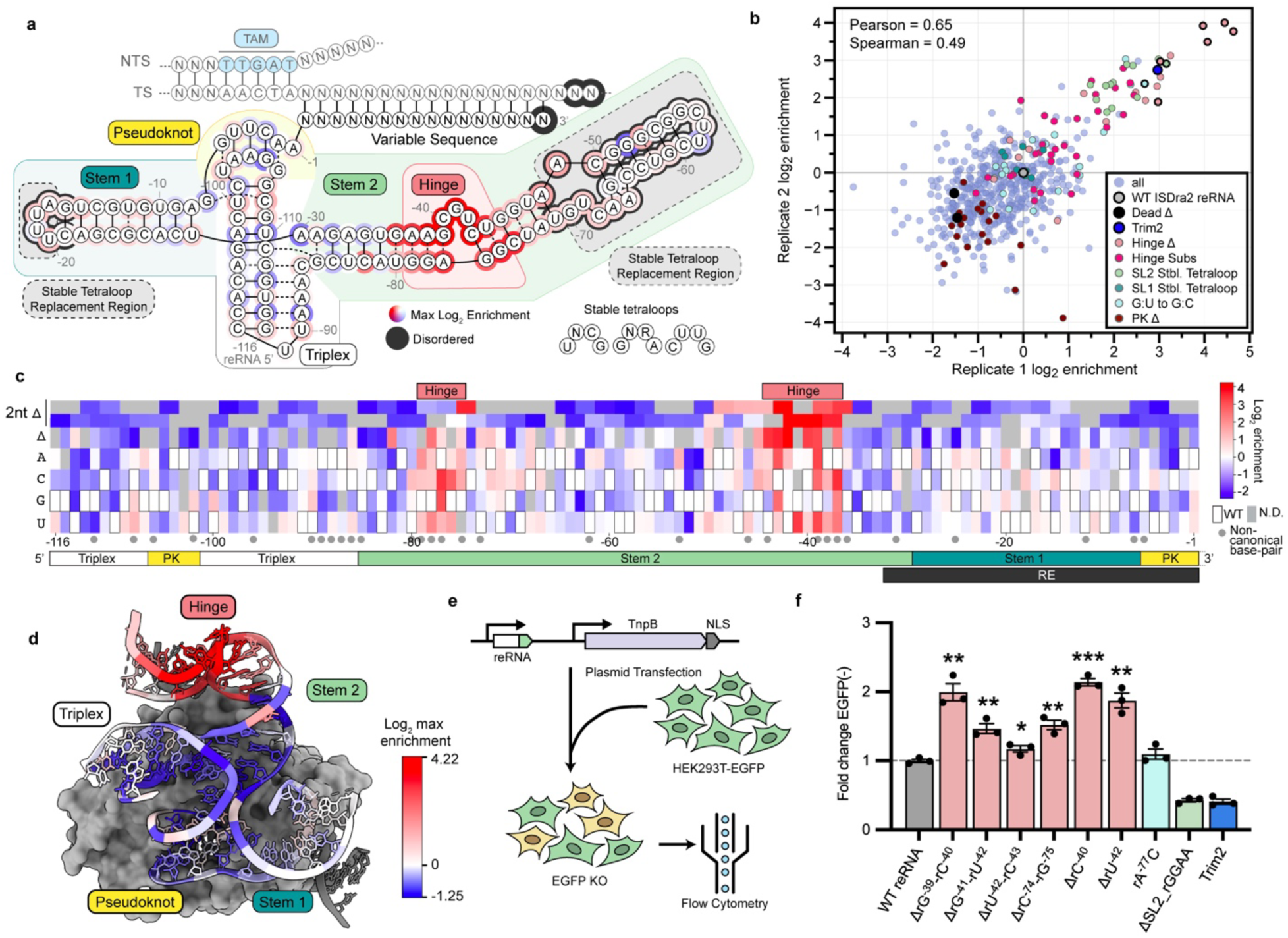
Profiling the TnpB reRNA mutational landscape reveals single nucleotide gain-of-function mutations. **a,** Schematic of the ISDra2 reRNA based on the RNP cryo-EM structure, adapted from Sasnauskas et al. 2023^6^. Dashed boxes indicate truncations replaced with stable tetraloops. Circles around bases are colored by maximum log2 enrichment scores for a substitution at each position, using the same color scale as in **c.** Outer gray-circled bases indicate regions not modeled in the ternary structure, labeled here as disordered. **b,** Log2 enrichment of reRNA variants across two experimental replicates. Data is normalized such that WT TnpB is at 0. **c,** Heatmap of enrichment scores for reRNA variants with single-nucleotide substitutions or single- and double-nucleotide deletions. WT positions are colored in white, and gray boxes denote no available data. **d,** Structure of TnpB ternary complex (PDB ID: 8EXA), with reRNA colored by positional maximum log2 enrichment scores **e,** Experimental workflow of EGFP knockout assay in HEK293T EGFP^+^ cells. EGFP^-^ cells were assessed by flow cytometry to compare editing activities of TnpB variants, as described in Methods. **f,** EGFP KO assay in HEK293T EGFP^+^ cells for highly enriched reRNA mutants identified with bold black outlines in **b**. Colors match the legend in **b**. Fold change of EGFP^-^ percent population for each variant compared to WT TnpB is shown. Data are presented as the mean ± s.e.m. (standard error of mean) from biological replicates (n = 3). Stars indicate a statistically significant increase in indel frequencies compared to WT ISDra2 reRNA as calculated using a two-sided unpaired Student’s t-test. (Significance: *, **, *** for p ≤ 0.05, 0.01, 0.001, respectively).

Upon mapping the reRNA mutational landscape, we found that inactivating truncations and deletions within the pseudoknot were depleted, as expected, under selective conditions (Fig. 2b). Stable tetraloop replacements in stem 2 were more highly enriched compared to those in stem 1, and Trim2 was one of the most highly enriched variants. Unexpectedly, we observed substitutions and deletions that were more enriched than both WT and the Trim2 variant. These activating mutations were concentrated around unpaired nucleotides rA^-40^-rU^-43^ within stem 2 (Fig. 2a,c). We refer to this region (rA^-37^-rU^-44^; rG^-75^-rG^-79^) as the “hinge” region of the reRNA, which appears to create a sharp bend in stem 2 preceding the disordered distal end^6^ (Fig. 2d). We also tested reRNA variants in HEK293T cells using an enhanced green fluorescent protein knockout (EGFP KO) assay^17^. When targeted to an EGFP transgene, reRNA variants with nucleotide deletions in the hinge region resulted in the greatest increase in EGFP KO compared to the WT reRNA, as assessed with flow cytometry (Fig. 2e-f). In contrast, reRNA variants with a truncation in stem 2, including Trim2, showed an unexpected decrease in EGFP KO efficiency compared to the WT reRNA, despite being highly enriched in the yeast selection (Fig. 2b, f).

Stem 2 was proposed by Sasnauskas *et al.* to act as a regulatory switch that controls the transition of the TnpB RNP into a cleavage-competent conformation upon DNA binding and heteroduplex formation^6^. This activation is driven by a conformational change in stem 2, where the formation of the RNA-DNA heteroduplex displaces the distal end of stem 2, leading to the release and activation of the RuvC domain. We speculate that activating hinge mutations enhance TnpB activity by increasing the flexibility of the distal end of stem 2, making it more prone to displacement. This could facilitate the release and activation of the RuvC domain. This aligns with the previous report that truncation of stem 2 also leads to dysregulated collateral ssDNA cleavage, independent of target DNA binding^6^. Variability in editing levels that we observe with stem 2 truncation variants may also support the importance of stem 2 in regulating reRNA-mediated TnpB activity (Fig. 2f). However, the precise mechanism underlying increased activity of reRNA variants remains unclear and requires further biochemical and structural investigation.

### Deep Mutational Scanning of TnpB Protein

We next mapped the fitness landscape of the ISDra2 TnpB protein with a library spanning all possible single amino acid substitutions and stop codons, alongside catalytically inactive (dead) and WT protein controls (Fig. 3). We collected data on 93% (7,611 of 8,140) of all possible substitutions across two biological replicates, which were highly reproducible with a Pearson correlation of 0.81 (Fig. 4a). We found that 3.7% of these substitutions were enriched by at least two-fold compared to WT TnpB (Figs. 3 and 4a).

**Figure 3.**
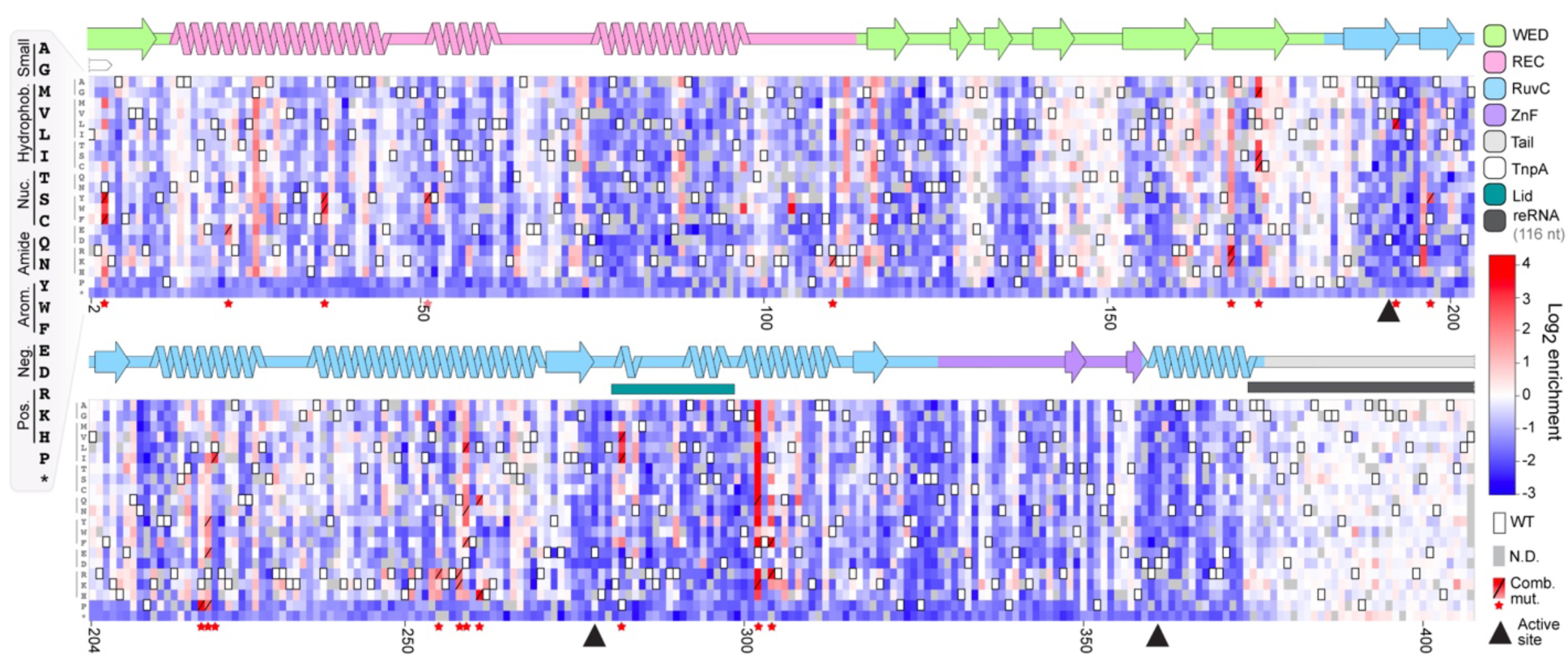
Deep mutational scanning of the TnpB protein identifies mutations that increase activity. Heatmap showing log2 enrichment scores of all single amino acid changes. TnpB secondary structures and domains are annotated above the heatmap. Outlined white boxes represent wild-type residues. Gray boxes denote positions with no available data. Boxes with a slash and a star at the x-axis represent a mutation that was included in the combinatorial variant library. Black triangles represent active site catalytic residues.

**Figure 4.**
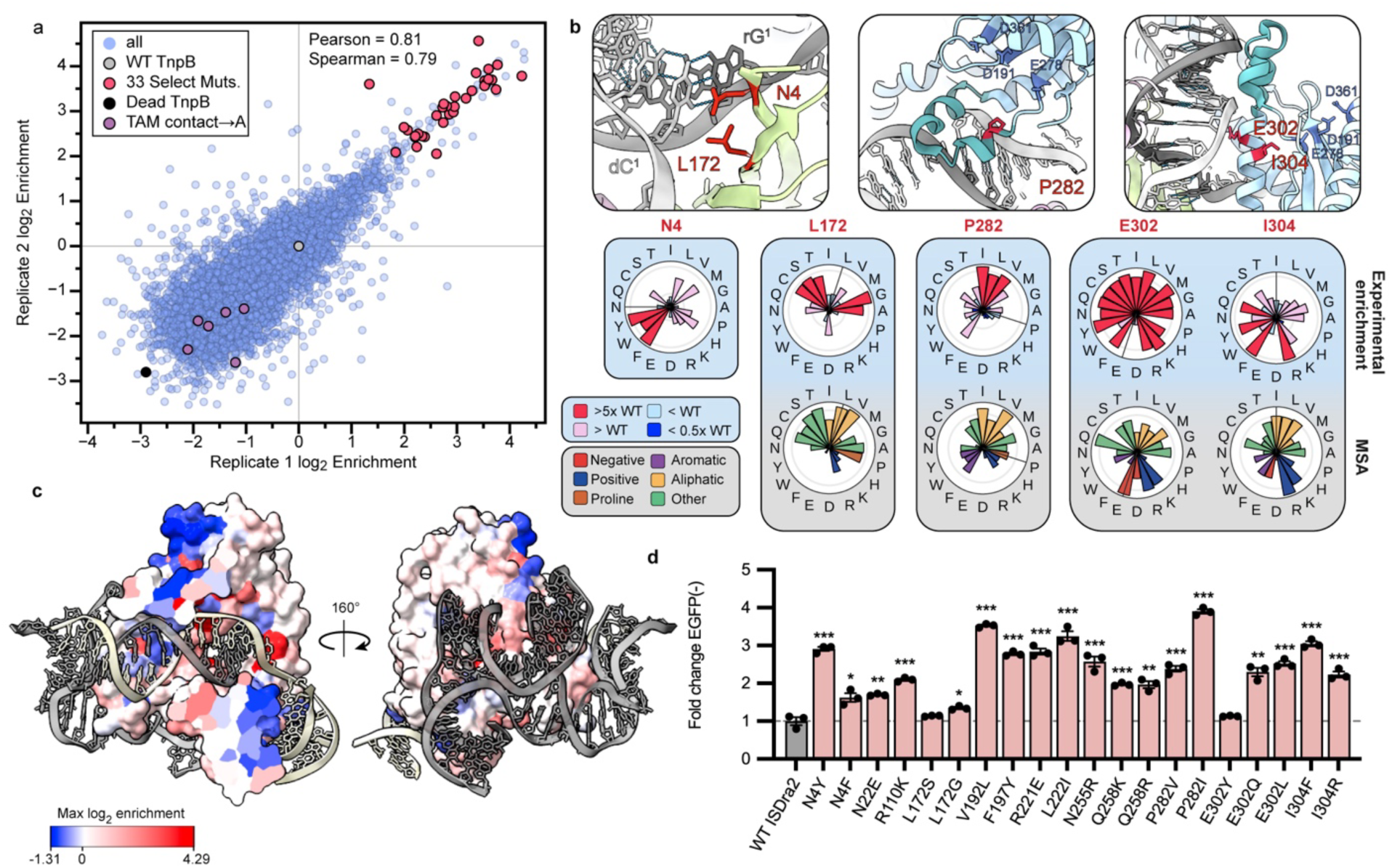
Activating mutations inform mechanistic insights and engineering. **a,** Log2 enrichment of single amino acid variants across two experimental replicates. Data is normalized such that WT TnpB is at zero. Thirty-three mutations were selected from the combinatorial library. Purple points correspond to alanine mutations at the TAM-interacting residues (Y52A, K76A, Q80A, T123A, S56A, F77A, N124A) shown to abrogate or reduce TnpB activity^7^. **b,** Close-up view of residues N4, P282, E302, and I304 in red in the ISDra2 TnpB ternary structure (PDB ID: 8EXA). RuvC catalytic residues are colored and labeled in dark blue. Site-wise amino acid enrichment and multiple-sequence alignment (MSA) conservation are displayed on radar plots below. N4 MSA data not shown due to low sequence conservation at the N-terminus. **c,** Max enrichment per residue mapped onto the surface of ISDra2 TnpB ternary structure, shown as a cross-section (left), allowing for visibility of the heteroduplex (PDB ID: 8EXA). The reRNA is shown in gray and DNA in light yellow. **d,** Activity of the highly enriched single amino acid mutants was assessed with the EGFP KO assay in HEK293T EGFP^+^ cells. Fold change of EGFP^-^ cells (percent of population) for each variant relative to WT TnpB is shown. Data are presented as the mean ± s.e.m. from biological replicates (n = 3). Stars indicate a statistically significant increase in indel frequencies compared to WT ISDra2 TnpB as calculated using a two-sided unpaired Student’s t-test. (Significance: *, **, *** for p ≤ 0.05, 0.01, 0.001, respectively).

Enrichment density varied across domains, with many mutations enriched in the RuvC and WED domains, most ZnF mutations depleted, and mutations in the unstructured C-terminal tail largely neutral (Fig. 3, Extended Data Fig. 2a). Stop codons causing truncations were depleted except for those at the C-terminus following residue 376, consistent with previous *in vitro* data showing that the C-terminal tail is dispensable for target cleavage^7^ (Extended Data Fig. 2b). Additionally, we observed depletion of alanine substitutions at residues important for recognizing the TAM, a 5’-TTGAT-3’ sequence which is essential for cleavage at the adjacent target sequence^2,3,7^ (Fig. 4a).

Positively charged amino acids were enriched over negatively charged residues within the vicinity of nucleic acids, particularly within the central channel, where the TAM-proximal end of the heteroduplex is accommodated^6^ (Extended Data Fig. 3). This is consistent with previous findings in CRISPR-Cas12 enzymes, where mutations to positively charged amino acids near the guide RNA-DNA heteroduplex, close to the protospacer adjacent motif (PAM), have been shown to increase activity and affect specificity^18,19^.

Similarly, specific WED-domain residues, N4 and L172, which stabilize the first TAM-proximal base pair of the heteroduplex^6^, were enriched for aromatic and small functional groups, respectively (Fig. 4b). We reason that the initial reRNA-target duplex formation may be enhanced by small hydrophobic and nucleophilic amino acids at position 172, and by π-stacking interactions between aromatic residues at N4 and the first heteroduplex nucleobases. The introduction of aromatic amino acids near the first base-pair of the RNA-DNA heteroduplex has also been associated with increased activity in AsCas12f^20,21^, providing evidence for a shared activation mechanism within an interaction conserved across TnpB and CRISPR-Cas12f endonucleases.

Within the positively charged central channel, E302 was highly enriched for substitutions to any amino acid that was not negatively charged (Fig. 4b). Positioned near the heteroduplex backbone, E302 might lead to electrostatic repulsion with the target strand (TS) phosphate backbone, potentially reducing RNP activity. We also identified other mutational hotspots, such as I304, where substitutions were enriched for residues with a range of physicochemical properties.

Unexpectedly, hydrophobic amino acids were enriched at position P282, which lies at the boundary of the lid subdomain that blocks the RuvC active site from accessing the TS^6^ (Figs. 3 and 4b). The lid subdomain forms non-sequence-specific contacts with the heteroduplex minor groove, which may aid in sensing heteroduplex formation prior to RuvC activation. We hypothesize that substituting the wild-type proline residue at this position with small, hydrophobic residues could increase the flexibility of the lid subdomain and accelerate the conformational change required to sense heteroduplex formation prior to TnpB activation.

Overall, 844 mutations from the DMS dataset were enriched over WT TnpB with a p-value <0.05 and were distributed across both the nucleic acid binding interface and the protein surface (Fig. 4c). We tested twenty highly enriched mutants in HEK293T cells with the EGFP KO assay. All reduced EGFP expression, with P282I resulting in a nearly four-fold reduction relative to WT TnpB (Fig. 4d).

To explore the generalizability of this protein DMS dataset, we transferred pairs of activating mutations to TnpB orthologs ISYmu1 and ISAba30^22^. These proteins share significant structural similarity with ISDra2 TnpB (pairwise TM-scores of 0.93 and 0.92 with ISYmu1 TnpB and ISAba30 TnpB, respectively), despite low sequence similarity (59% with ISYmu1 and 48% with ISAba30)^23,24^. We designed two ISYmu1 TnpB variants (H4Y/V305R and L167G/V305R) and one ISAba30 TnpB (L4Y/V272I) variant by introducing pairs of analogous activating mutations from ISDra2 TnpB (N4Y/I304R; L172G/I304R; and N4Y/P282I, respectively) (Extended Data Fig. 4a). We observed increased colony reversion with all three variants compared to their WT orthologs in the yeast cleavage assay (Extended Data Fig. 4b-c). These results demonstrate that these activation mechanisms are generalizable beyond ISDra2 TnpB, underscoring the utility of our mutational dataset for informing further engineering and characterization of diverse systems.

### Combinatorial mutations enhance TnpB activity

To explore increases in TnpB editing activity through mutation combinations, we selected 33 highly enriched single amino acid mutations covering 19 positions across TnpB (Fig. 3). Using nicking mutagenesis, we generated a library of ∼5×10³ variants with an average of ∼5 of the 33 possible mutations per variant^25,26^ (Extended Data Fig. 5a). This combinatorial variant library underwent selection in two reporter yeast strains with different target sequences (Fig. 5a, Extended Data Fig. 5b). We observed the greatest increase in enrichment in variants with 4-5 mutations on average, while depleted variants had more variable mutation numbers (Extended Data Fig. 5c). The expression levels of seven highly active TnpB variants were found to be similar to wild-type by Western blot, consistent with a change in enzymatic activity rather than protein abundance (Extended Data Fig. 6).

**Figure 5.**
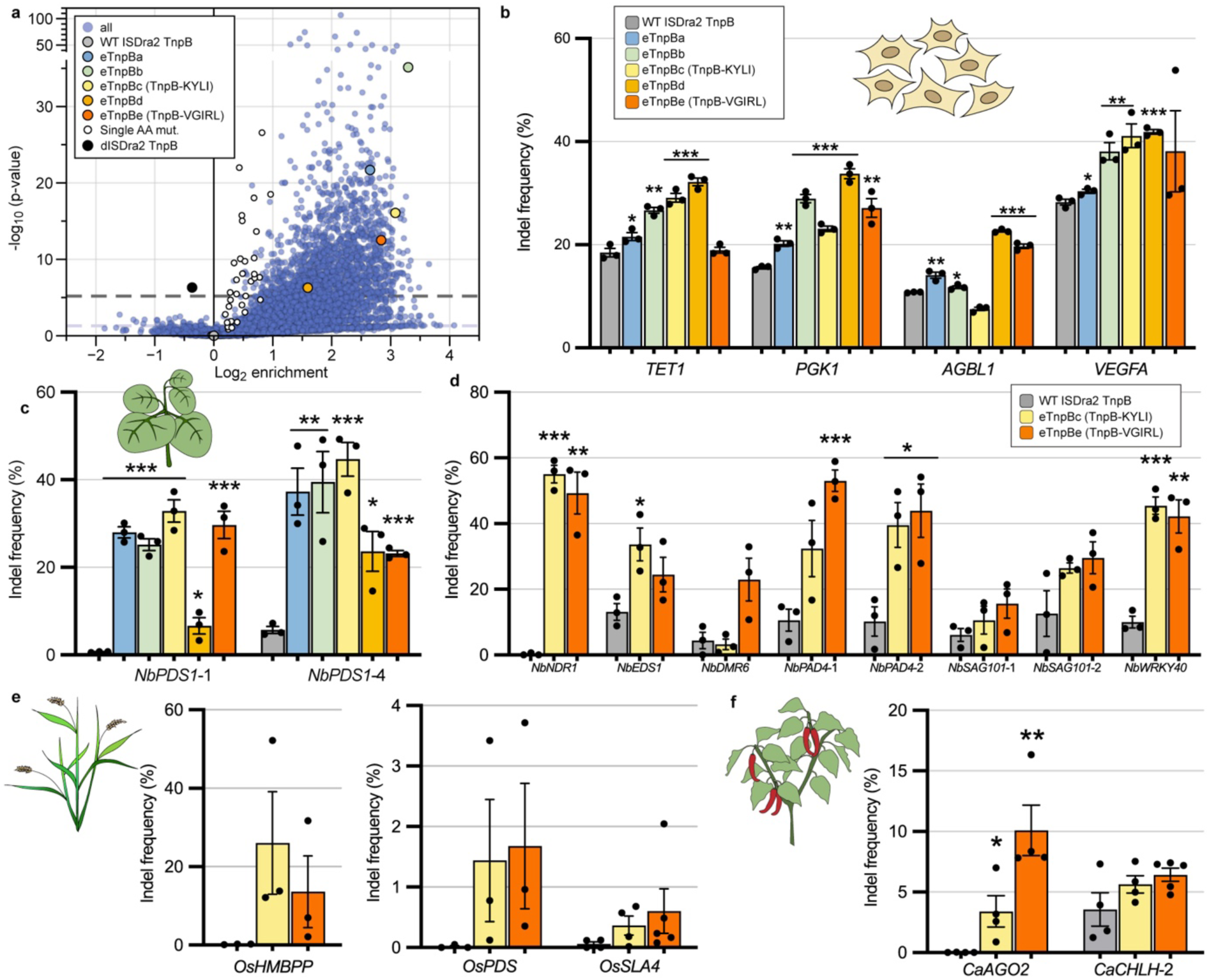
Enhanced TnpB variants engineered by combining high-activity mutations. **a,** Volcano plot depicting combinatorial variant average enrichment and statistical significance, after selection in a reporter yeast strain. Individual barcodes were used to calculate significance with a two-sided Mann-Whitney U-test. **b,** Indel frequency of five combinatorial variants at four genomic loci in HEK293T cells. Plasmids carrying TnpB variants were delivered via transfection, and genomic DNA was sequenced as described in methods. **c,** Indel frequency of TnpB variants targeting two sites in *PDS1* in *N. benthamiana*. Legend, same as in **b**. **d,** Indel frequency of WT ISDra2, TnpB-KYLI, and TnpB-VGIRL targeting eight sites in *N. benthamiana*. **e,** Indel frequencies of ISDra2 TnpB variants targeting three sites in rice callus tissue. **f,** Indel frequency of TnpB variants targeting two sites in pepper. TnpB variants were delivered by agroinfiltration, and indel frequencies were quantified from harvested tissue as described in the methods for **c-f**. Data are plotted as the mean and s.e.m. from biological replicates (n≥3) in **b-f**. Stars indicate a statistically significant increase in indel frequencies compared to WT ISDra2 TnpB as calculated using a two-sided unpaired Student’s t-test. (Significance: *, **, *** for p ≤ 0.05, 0.01, 0.001, respectively) in **b-f**.

To assess the genome editing activity of TnpB combinatorial variants, we selected five highly active variants (eTnpBa–eTnpBe) for testing at five genomic loci in human cells (Fig. 5b, Extended Data Fig. 7a). HEK293T cells were transfected with plasmids encoding each TnpB protein variant targeting endogenous loci, and indel (insertion/deletion) frequencies were assessed four days post-transfection. Compared to WT TnpB, all five variants demonstrated higher indel formation frequencies across all target sites, except for eTnpBc (N4Y/R110K/V192L/L222I) at the *AGBL1* locus. Variant eTnpBd (R110K/P282V/E302Q) achieved the highest overall indel frequencies across multiple loci (23-42%), surpassing both wild-type ISDra2 and ISYmu1 TnpB (11-29% and 6-30%, respectively) (Fig. 5b, Extended Data Fig. 7a).

To evaluate off-target activity in HEK293T cells, we identified six genomic sites with 4– 6 mismatches to the target-complementary reRNA sequence with Cas-OFFinder^27^. All variants exhibited increased off-target indel frequencies compared to wild-type ISDra2 and ISYmu1 on at least two sites, with up to 6% off-target indel frequencies observed with eTnpBc at *TET1* off-target site 1 (Extended Data Fig. 7b). Generally, lower indel frequencies occurred at off-target sites with more TAM-proximal mismatches preceding the 12th nucleotide, consistent with published data indicating that TAM-distal mismatches are more well-tolerated by WT TnpB^6,28^. While increases in on-target editing were consistently accompanied by increases in off-target indel frequencies, eTnpBe (L172G/V192L/L222I/P282V/I304R) was associated with the lowest off-target activity (<2% at all sites) of the variants.

We investigated whether genome editing activity could be further enhanced by combining highly active TnpB protein with reRNA variants (Extended Data Fig. 7c). Using the EGFP-KO assay, we tested five TnpB protein mutants paired with one of two reRNA mutants (ΔU^-42^ or ΔC^-74^-G^-75^), each with deletions on opposite strands of the hinge region. However, many of these pairings with the reRNA mutants did not result in substantial additive improvements in editing, compared to pairings with the WT reRNA. Instead, combining the highly active eTnpBd variant with either hinge deletion variant resulted in reduced genome editing (15-24% EGFP^-^ cells) compared to the wild-type reRNA (44.5% EGFP^-^ cells). One possible explanation is that certain protein-reRNA variant combinations destabilize the TnpB RNP beyond a critical free energy threshold, disrupting RNP assembly or dsDNA targeting^29^. Further studies are needed to understand how protein and reRNA mutations interact to influence RNP stability and function. Systematic combinatorial testing of reRNA and protein variants may reveal more optimal pairings that enhance RNP stability and activity.

### Enhanced TnpB variants for genome editing in plants

Precise genome editing offers major advantages over traditional breeding in identifying and developing novel crop traits, but is still limited by efficient delivery of gene editing components and the low-throughput of plant tissue culture^30,31^. Viral vectors have shown promise in delivering gene editing reagents to induce heritable germline edits across various species; however, their cargo capacity remains a significant limitation for delivering standard CRISPR-Cas enzymes^32,33^. TnpB, due to its compact size, is well-suited to overcome this barrier. Multiple TnpB orthologs, including ISDra2, have shown potential for genome editing in plants, but wild-type editing efficiencies remain low^34–36^.

To assess their utility for plant genome editing, we tested eTnpBa-eTnpBe in the model dicot *Nicotiana benthamiana*. TnpB variants and reRNA targeting three sites within *NbPDS1* (*Phytoene desaturase*) were delivered into *N. benthamiana* leaves by agroinfiltration, and indel frequencies were assessed within the infiltrated leaf tissue. All five variants exhibited increased editing activity at *NbPDS1*-1 and *NbPDS1*-4 sites (Fig. 5c). At both *NbPDS1*-1 and *NbPDS1*-4, eTnpBc demonstrated the highest editing efficiencies of (33% and 45%) compared to WT TnpB (<1% and 6%), with all variants demonstrating editing levels between roughly four to forty-fold higher than WT levels. At the *NbPDS1*-2 site, an increased indel frequency was observed for eTnpBe (8% vs. 2% WT TnpB) (Extended Data Fig. 8a). In contrast, we did not observe substantial increases in editing with the reRNA and single amino acid substitution variants compared to WT ISDra2 at *NbPDS1*-1 and *NbPDS1*-2 (Extended Data Fig. 8b).

Based on their activity across multiple genomic sites in HEK293Ts and *N. benthamiana*, we selected eTnpBc and eTnpBe, hereafter referred to as TnpB-KYLI (R110**K**/N4**Y**/V192**L**/L222**I**) and TnpB-VGIRL (P282**V**/L172**G**/L222**I**/I304**R**/V192**L**), for assessment at eight additional genomic target sites in *N. benthamiana* (Fig. 5d). Except for TnpB-KYLI at the *NbDMR6* site, both TnpB-KYLI and TnpB-VGIRL exhibited increased editing activity compared to WT ISDra2 at all sites, with TnpB-KYLI and TnpB-VGIRL reaching over fifty-fold increases in indel frequencies (54% and 50%) compared to WT ISDra2 (<1%) at *NbNDR1*. At the *NbDMR6* site, TnpB-VGIRL exhibited an increased indel frequency (23%) relative to WT ISDra2 (4%). Furthermore, editing levels of TnpB-KYLI and TnpB-VGIRL at off-target sites predicted by Cas-OFFinder^27^ were comparable to or lower than those observed for the WT TnpB, indicating the engineered variants maintain target specificity in *N. benthamiana* (Extended Data Fig. 9a).

We compared the editing efficiencies of TnpB-KYLI and TnpB-VGIRL with WT ISDra2 and ISYmu1 TnpB, as well as other recently engineered small RNA-guided endonucleases, such as AsCas12f-HKRA^21^ and NovaIscB^37^, which also offer advantages for viral delivery where cargo size is limited. Among these small RNA-guided endonucleases, TnpB-VGIRL and TnpB-KYLI showed the highest editing levels at all three target sites in *N. benthamiana*, with TnpB-VGIRL nearly matching the indel frequencies observed with Cas9 at *NbWRKY40* (38% and 39% for TnpB-VGIRL and Cas9, respectively) (Extended Data Fig. 9b). In comparison, the editing levels of AsCas12f-HKRA, NovaIscB, and ISYmu1 were consistently lower (<9%) or undetectable.

We speculate that the relatively low editing efficiencies of AsCas12f-HKRA and NovaIscB in *N. benthamiana* may be due to differences in the strategies used to engineer these RNA-guided endonucleases for increased activity. While AsCas12f-HKRA and NovaIscB were optimized using HEK293T cell-based assays or *in vitro* cleavage assays at or above 37°C^21,37^, our selections were performed in yeast, which grow at 30°C, and may have enriched for variants with greater activity at lower temperatures optimal for plant growth (23-28°C, see Methods).

Finally, we investigated the ability of the variants to edit in the agriculturally important monocot and dicot crop species, rice (*Oryza sativa*) and pepper (*Capsicum annuum*). To assess the potential for generating stable transgenic lines, we measured editing in rice callus transformed with *Agrobacterium* carrying TnpB-KYLI, TnpB-VGIRL, and WT TnpB targeting three genomic loci. In rice calli, TnpB-KYLI and TnpB-VGIRL showed higher editing levels than WT TnpB at all three sites, with up to 26% and 14% indel frequencies observed for TnpB-KYLI and TnpB-VGIRL at *OsHMBPP* (Fig. 5e). Although rice callus transformation methods are well-established and WT ISDra2 TnpB editing has been reported in rice, effective delivery of genome editing components into pepper remains limited^38–40^. To further demonstrate editing activity of TnpB-KYLI and TnpB-VGIRL in a non-model crop, we delivered these variants and reRNA targeting five genomic sites into pepper leaves by *Agrobacterium* infiltration. TnpB-KYLI and TnpB-VGIRL demonstrated consistently higher editing compared to WT TnpB at all five sites, with up to 10% editing with TnpB-VGIRL at *CaAGO2*, compared to <1% editing with WT TnpB (Fig. 5f, Extended Data Fig. 9c). Overall, these findings highlight the potential of TnpB-KYLI and TnpB-VGIRL for high-efficiency editing in both crops, and further optimization of delivery strategies may enable higher editing levels.

## Conclusions

In the *Deinococcus radiodurans* genome, ISDra2 TnpB is encoded alongside the HUH superfamily TnpA transposase, with both relying on overlapping sequences essential for transposition and endonuclease activity^2,41^. The reRNA stem 1 sequence overlaps with the imperfect hairpin in the transposon right end (RE), which is required for TnpA recognition and excision^6,41,42^. Notably, in our DMS datasets, stem 1 exhibited a high degree of mutational tolerance, as did the few protein residues within the vicinity of stem 1, supporting the hypothesis that TnpB protein and reRNA have co-evolved with TnpA and the transposon^43^ (Extended Data Figs. 3a and 10).

The selective balance of transposon maintenance, propagation, and effects on host fitness may constrain TnpB nuclease activity in its native setting^12,44,45^. By profiling TnpB-mediated on-target cleavage outside the context of transposition, we identified many activating mutations across the RNP, highlighting the rugged nature of the mutational landscape^7,46^. The frequency of activating and neutral mutations in TnpB is an outlier compared to standard models of protein evolution and other mutational studies^29,47–49^. This aligns with the hypothesis that TnpB exhibits pervasive evolutionary flexibility, having been exapted for diverse biological processes across multiple clades of life^1,43,50,51^. Additionally, the prevalence of evolutionarily-accessible activating mutations may suggest TnpB endonuclease activity is under negative selective pressure in the transposon context.

This work presents the first comprehensive sequence-function landscapes for both the protein and RNA scaffold of an RNA-guided endonuclease. Comprehensive reRNA mutagenesis uncovered an unexpected mutational hotspot in stem 2, and offers an alternative approach to iterative reRNA and gRNA engineering through truncations and G:U swaps to optimize gene editing activity^6,21,52^. Mutational scanning of the TnpB protein not only reproduced published findings on ISDra2 TnpB point mutants^7,46^, but also captured new activating mutations that increased on-target cleavage activity. We further demonstrate that activating mutations can be combined to enhance genome editing activity in HEK293T cells, *N. benthamiana,* rice, and pepper, and we present TnpB-KYLI and TnpB-VGIRL as highly active variants. These small, engineered TnpB variants offer a promising solution for overcoming current cargo restrictions that limit the delivery of transiently-expressed genome editors in plants^31^. Further exploration in plants with alternative delivery methods and broadening of the 5’-TTGAT-3’ TAM recognition motif will increase the utility of TnpB variants for genome editing applications^53^.

Overall, these comprehensive mutagenesis libraries provide molecular insights into nucleic acid binding, activation, and cleavage by TnpB, mapping both mutational constraints and activating mutations across the ribonucleoprotein. Further biochemical and epistatic studies may help elucidate mechanisms of activating mutations in TnpB and related endonucleases. In addition to laying the groundwork for further engineering, we hope our findings will provide insights into TnpB’s evolution and function within insertion sequences.

## Acknowledgements

We thank Joseph Luis Rivera for cultivating the *N. benthamiana* plants used in this study. We thank Noam Prywes (University of California, Berkeley, CA) for advice on library design and general guidance on deep mutational scanning and data analysis. We thank Luke Oltrogge (University of California, Berkeley, CA) for key assistance with data analysis and sequencing. We thank Kai Chen (University of California, Berkeley, CA) for providing the HEK293T-EGFP cell line used for EGFP knock-out assays. We thank John Desmarais (CSHL), Brady Cress (University of California, Berkeley, CA), Amy Eggers (Seek Labs), and Joshua Cofsky (Harvard University) for establishing protocols and providing components of expression vectors that we used in the yeast cleavage assay. Antiserum against phosphoglycerate kinase (PGK) was generously provided by Dr. Jeremy Thorner (University of California, Berkeley, CA)^54^. We thank Netra Krishnappa (University of California, Berkeley, CA) for assistance in running NGS samples, and Julia Turnšek (University of California, Berkeley, CA) for help with preliminary protein purification. We additionally thank Owen Tuck (University of California, Berkeley, CA) for insights into the protein DMS dataset and experimental design, and Maria Lukarska (University of California, Berkeley, CA) for reviewing the manuscript.

## Author Contributions

BWT, RFW, DFS, and JAD conceptualized the project. DFS and JAD supervised the study. BWT and RFW designed and conducted the experiments with assistance from RVT and BTD. RFW and BWT analyzed the data. BWT, RFW, RVT, and BTD collected data on individual TnpB variants in yeast, HEK293Ts, and *N. benthamiana*. JER, UN, EDG, FW, VG, and collected editing data in *N. benthamiana* and additional plant species. CIT collected data on engineered TnpB ortholog activity in yeast. BWT, RFW, and JT developed the yeast assay. SPD-K, UN, BWT, and RFW collected editing data in pepper. M-JC, GA, BWT, and RFW collected editing data in rice. BWT, RFW, JP, DFS, and JAD wrote the manuscript with input from all authors. All authors reviewed the manuscript and approved the final version.

## Extended data

**Extended Data Figure 1.**
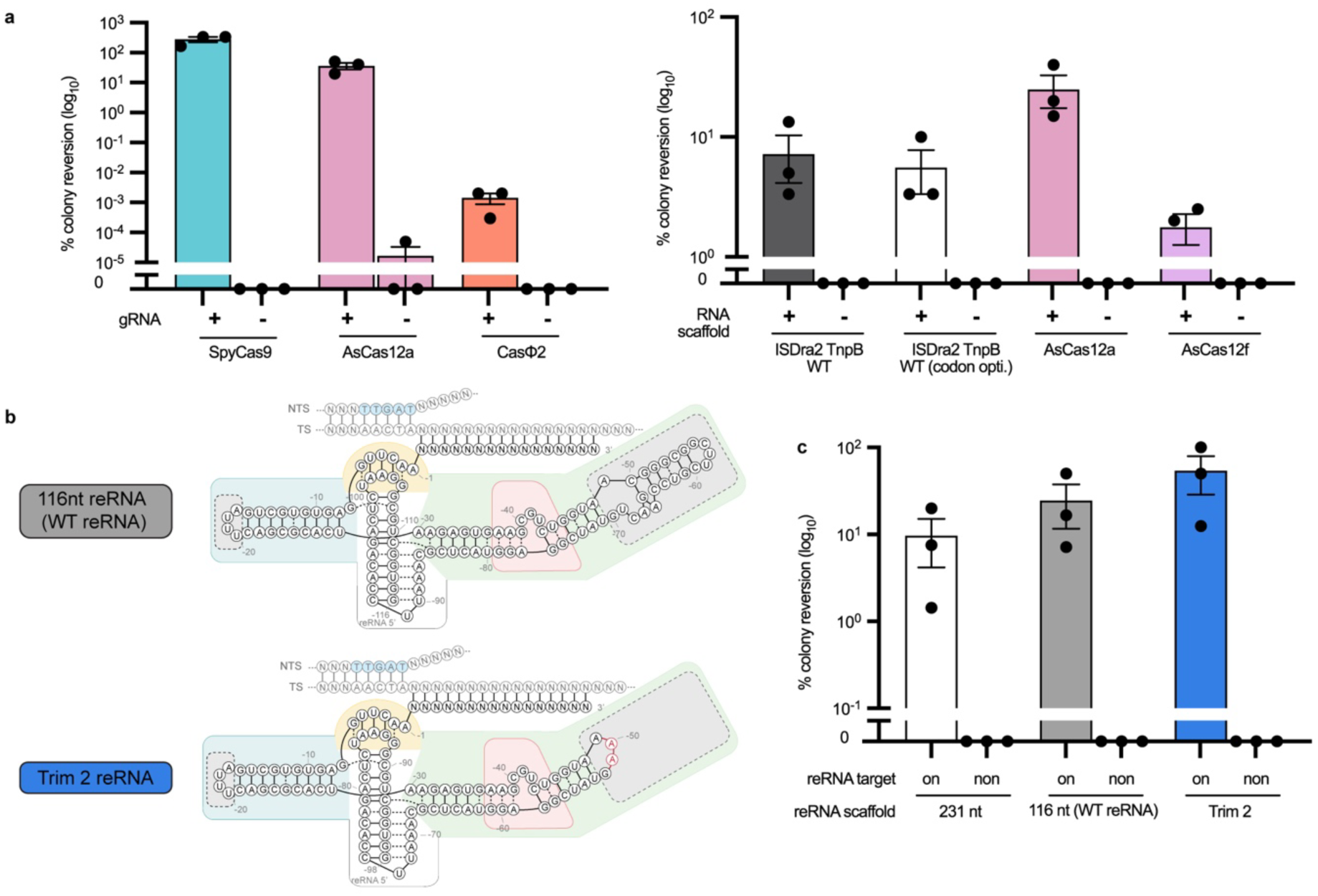
*In vivo* yeast cleavage assay captures a range of RNA-guided endonuclease activity across TnpB, Cas9, and Cas12 endonucleases. **a,** Assessment of Cas9, Cas12, and ISDra2 TnpB RNA-guided endonuclease activity in yeast cleavage assays, with (+) or without (-) the gRNA or reRNA. ISDra2 TnpB protein was expressed from either the non-codon-optimized open reading frame from *D. radiodurans* R1 (GenBank AE000513.1), or TnpB was codon optimized for high frequency codon usage between *H. sapiens* and *S. cerevisiae.* Data are plotted as the mean and s.e.m. (standard error of mean) from technical triplicate titer plating measurements (n=3). **b,** Schematic representation of the ISDra2 Trim2 reRNA variant (red rArA bases replacing ΔrC^-50^-rU^-69^). Color scheme corresponds to Fig. 2a. **c,** ISDra2 TnpB endonuclease activity in yeast with various reRNA scaffold lengths, including 231 nts, 116 nts, and the reported Trim2 reRNA variant. TnpB and reRNA were expressed in a yeast strain with an reRNA-complementary target site (on), or in a yeast strain with a non-complementary target site (non). Data are plotted as the mean and s.e.m. from technical triplicate titer plating measurements (n=3).

**Extended Data Figure 2.**
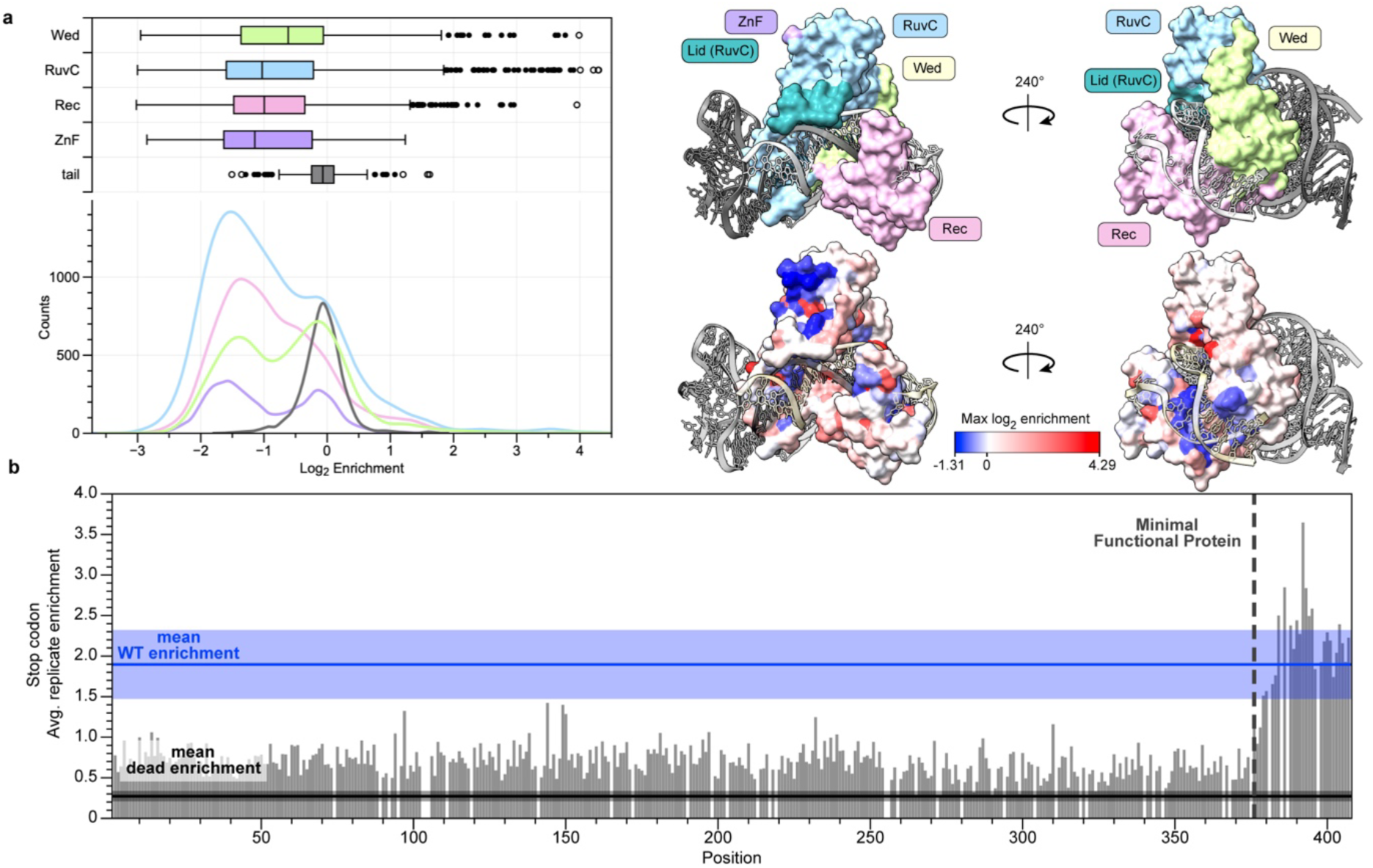
Distribution of enriched amino acid substitutions varies by TnpB domain. **a,** (Left) Histogram and box plots of the enrichment values for all protein DMS library mutations, separated by ISDra2 TnpB domain. (Right) ISDra2 domains (top right) and max enrichment per amino acid residue (bottom right) mapped onto the surface of ISDra2 TnpB ternary structure (PDB ID: 8EXA). Data are plotted as median+IQR. Outliers (Q1-1.5 IQR or Q3+1.5 IQR) drawn as circles and extreme outliers (Q1-3 IQR or Q3+3 IQR) are drawn as open circles **b,** Enrichment scores averaged for all stop codon mutations plotted across the length of the protein. Enrichment scores are not normalized to WT. The dashed line indicates position 376, marking the C-terminus of the minimal active TnpB truncation variant (Δ376–408) previously identified^7^.

**Extended Data Figure 3.**
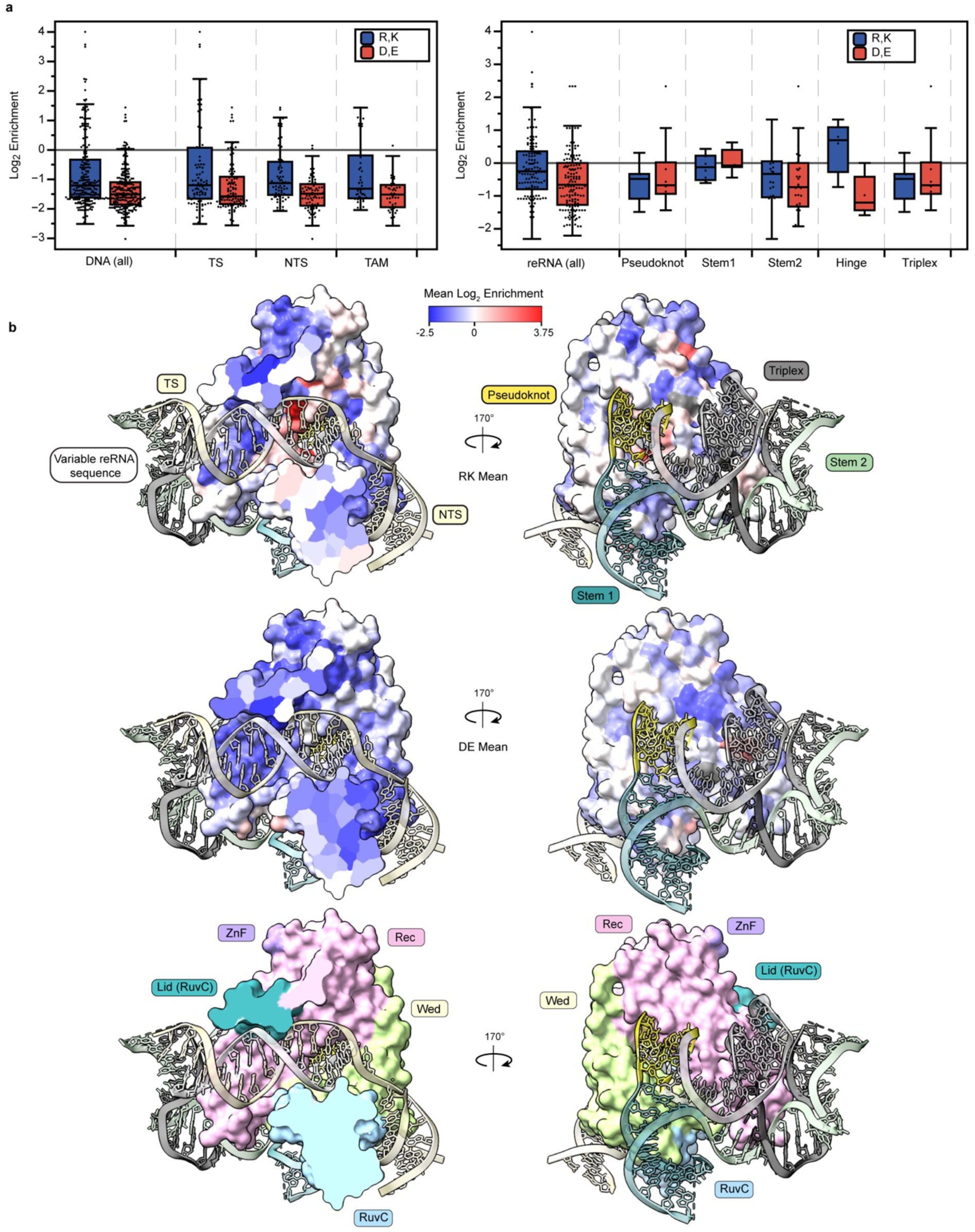
Positively charged amino acids are enriched near nucleic acid contacts. **a,** Enrichment of positively (R, K) and negatively (D, E) charged amino acid substitutions for residues proximal to nucleic acid. To assess the impact of amino acid substitutions near nucleic acids, we defined proximal residues as those with Cα atoms within 8Å from nucleic acid atoms. This cutoff ensured inclusion of previously identified direct interactions and potential contacts by R/K/D/E mutations. Data are plotted as median + IQR. **b,** The average enrichment for substitutions to positively charged (R, K) or negatively charged (D, E) amino acids was calculated at each position. If the WT residue was already R/K/D/E, its enrichment value was included in the average as zero. Enrichment values were mapped onto the ISDra2 TnpB cryo-EM ternary structure^6^, with additional surface coloring by domain to help orient the reader to the structural context.

**Extended Data Figure 4.**
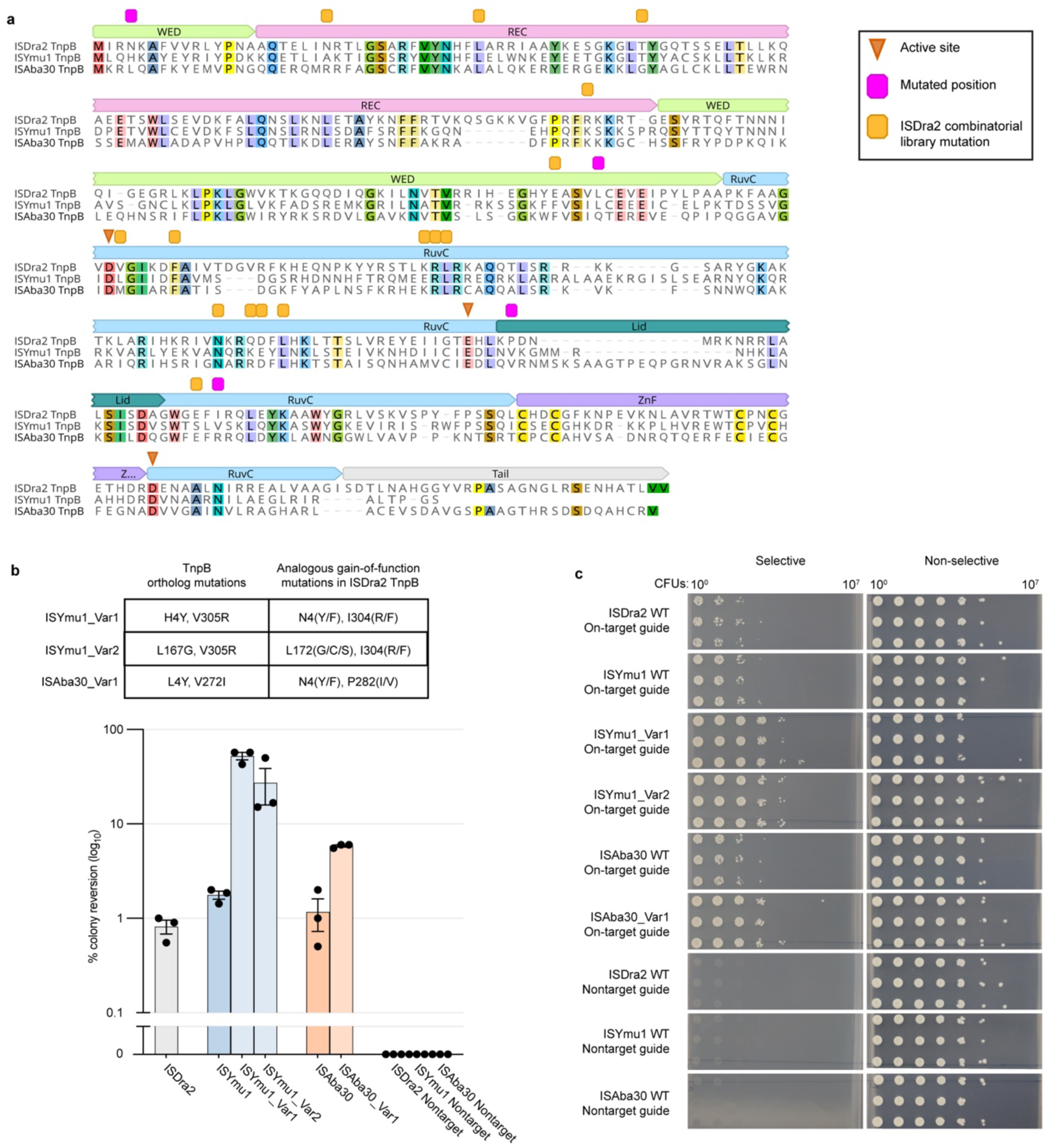
Activating mutations found for ISDra2 TnpB are transferable to TnpB orthologs. **a,** Multiple structural alignment of ISDra2, ISYmu1, and ISAba30 TnpB. **b-c,** Activity of ortholog mutants was assessed by percent colony reversion with the yeast cleavage and compared to WT orthologs and negative non-complementary reRNA-target controls. **b,** Data represent mean ± s.e.m. (n = 3 technical plating replicates 8 hours post-induction). **c,** Titer plates (**b**) where each of the protein variants have been expressed in *S. cerevisiae* for 8 hours and titer plated on selective (-adenine) and nonselective (+adenine) media. Technical pipetting replicates from titer plates are shown.

**Extended Data Figure 5.**
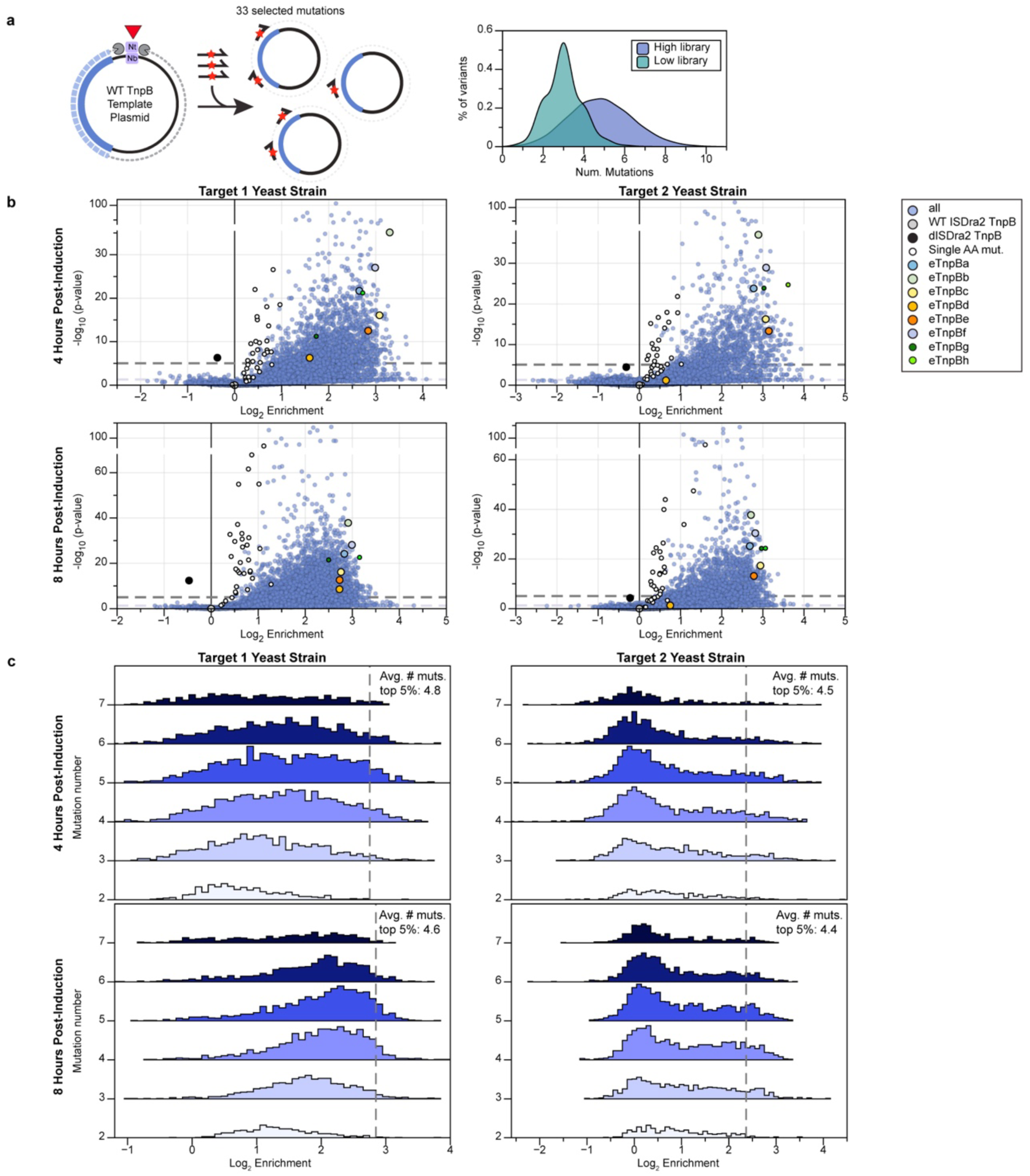
Combinatorial library construction, experimental enrichment, and distribution of mutation number. **a,** Schematic of library construction using pooled nicking mutagenesis. Plasmid was digested for ssDNA template, and 33 selected amino acid mutations at 19 positions were introduced on ssDNA oligos as described in methods. Two libraries with low and high mutation frequencies were combined, with an average of 3 and 5 mutations, respectively. **b,** Volcano plots of variant enrichment and statistical significance in orthogonal reporter yeast strains with different target sites. Enrichments are shown from 4 hours and 8 hours post-induction. Enrichment is calculated by averaging two biological replicates. Significance was calculated from individual barcode enrichments per variant relative to wild-type (two-sided Mann-Whitney U-test). **c,** Enrichment score distributions separated by variant mutation number for experimental conditions in **b.**

**Extended Data Figure 6.**
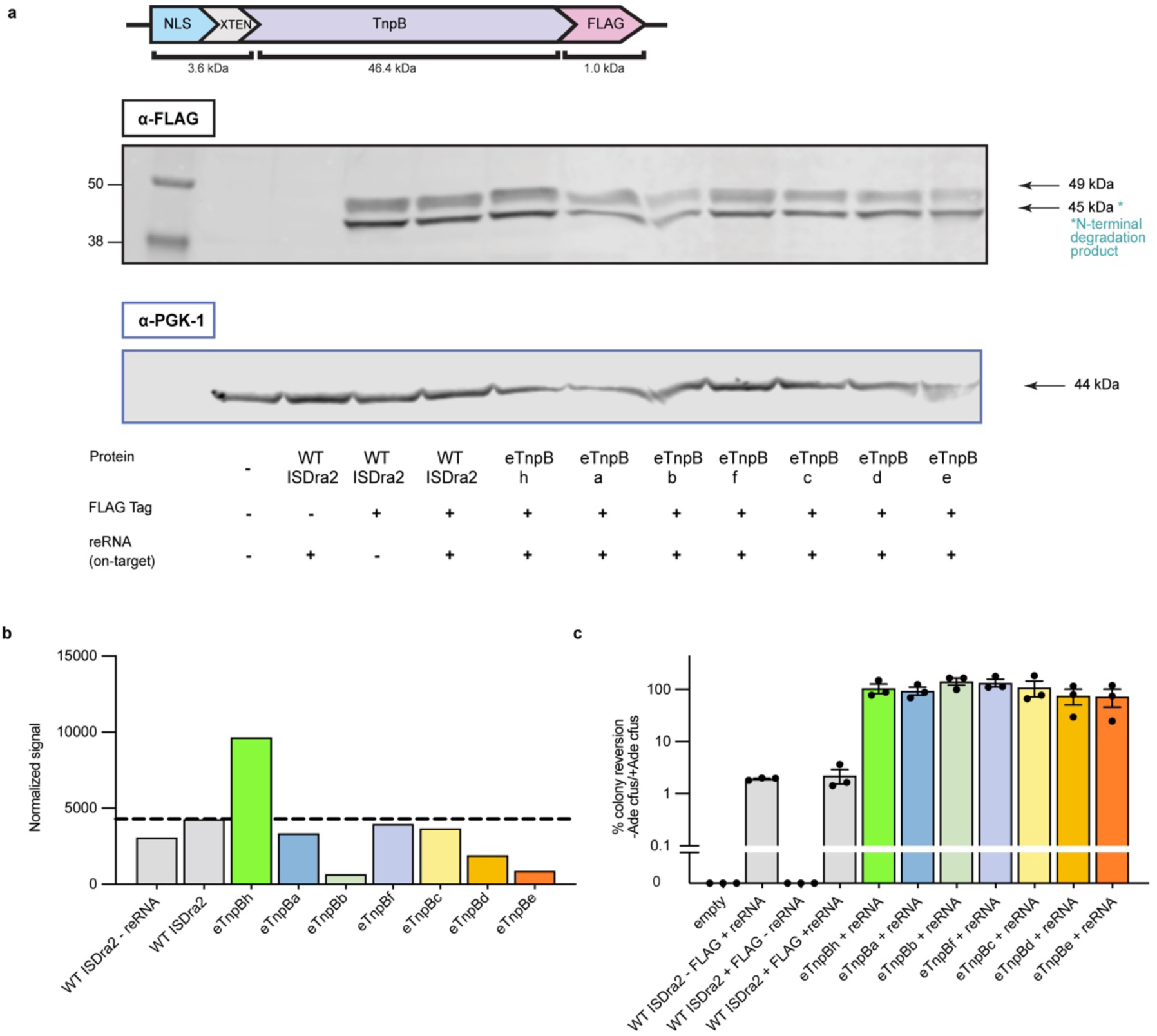
Western blots showing that expression levels of enhanced TnpB variants do not increase in *S. cerevisiae*. **a,** (Top) Construct design for expression of ISDra2 TnpB WT protein and variants with an NLS and FLAG tag in yeast. (Bottom) Western blot from yeast lysate with anti-FLAG antibody, and with an anti-PGK1 antibody as a loading control. **b,** Quantification of signal from Western blot for each variant **c,** Activity of each variant was assessed by colony reversion in the yeast cleavage assay. Data represent mean ± s.e.m. (n = 3 technical plating replicates).

**Extended Data Figure 7.**
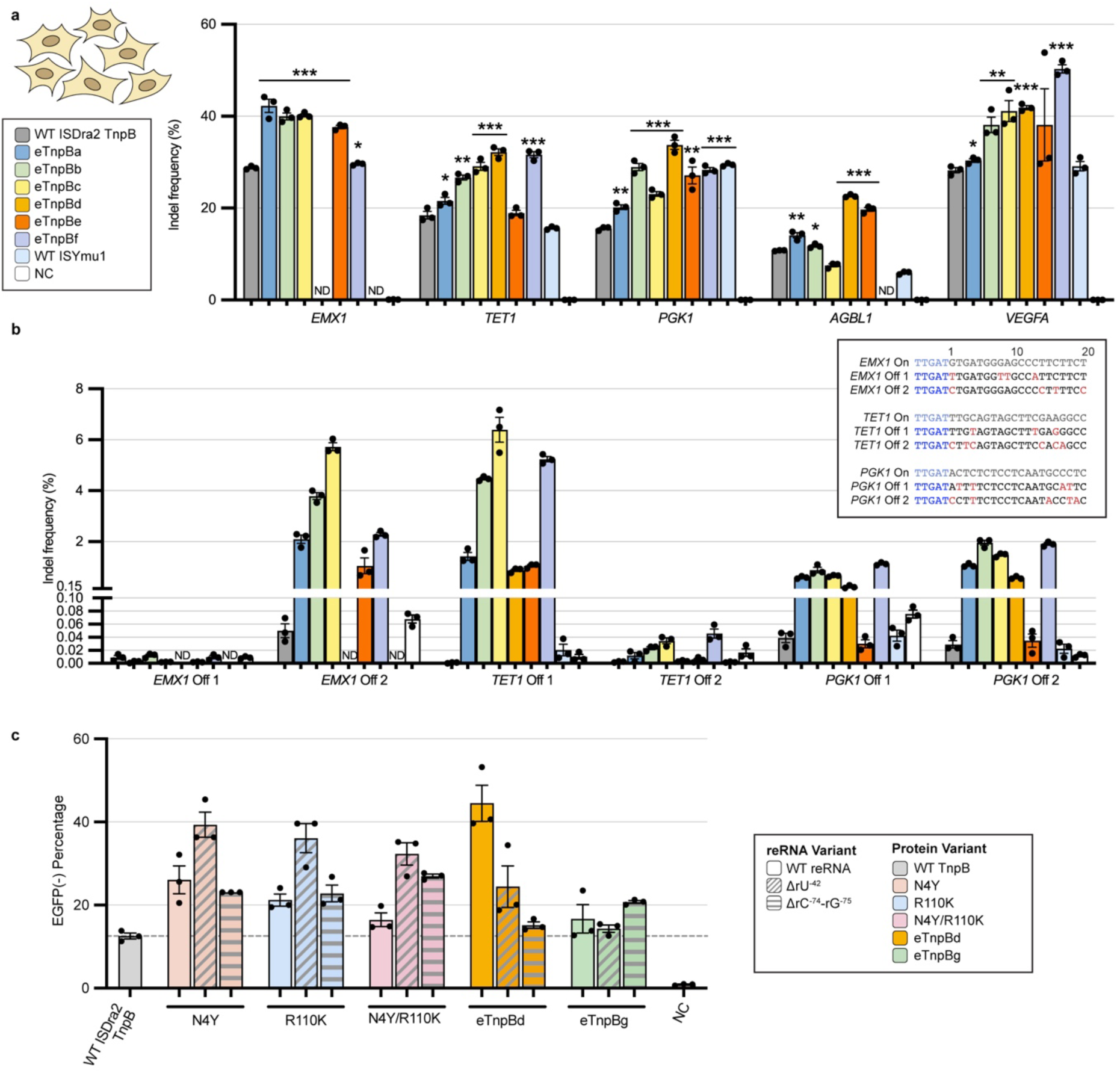
Assessment of combinatorial TnpB variant off and on-target editing, with reRNA mutants in HEK293Ts. **a,** Indel frequency of six combinatorial variants at genomic loci in HEK293T cells, with WT ISDra2 TnpB, WT ISYmu1 TnpB, and no plasmid (NC) controls. Indel frequencies for eTnpBa-eTnpBe and WT TnpB at *TET1*, *PGK1*, *AGBL1*, and *VEGFA* are also represented in Fig. 5b. Data are plotted as the mean and s.e.m. from biological replicates (n=3). ND indicates no data. Stars indicate a statistically significant increase in indel frequencies compared to WT ISDra2 TnpB as calculated using a two-sided unpaired Student’s t-test. (Significance: *, **, *** for p ≤ 0.05, 0.01, 0.001, respectively). **b,** Indel frequency of WT and combinatorial variant TnpBs at off-target sites identified by Cas-OFFinder, with 4–6 mismatches to three on-target sites. Sample order and color scheme match **a.** Off-target sequences (non-target strand) are listed, with TAM in blue and reRNA-target mismatches in red. Data are plotted as the mean and s.e.m. from biological replicates (n=3). **c,** Pairs of reRNA deletion and ISDra2 TnpB protein mutants were tested with the EGFP KO assay, where EGFP-negative cells were measured by flow cytometry seven days after transfection. Data are presented as the mean ± s.e.m. from biological replicates (n = 3).

**Extended Data Figure 8.**
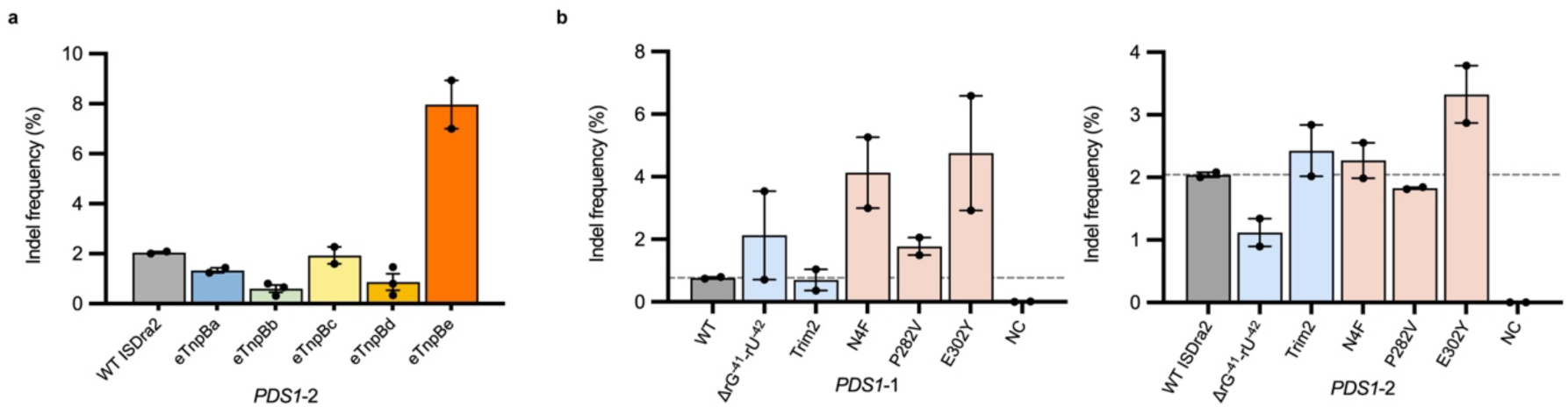
TnpB protein and reRNA mutants enable increases in TnpB-mediated indel frequencies in *N. benthamiana.* **a,** Indel frequencies of TnpB combinatorial protein mutants at *PDS1*-2 in *N. benthamiana*. **b,** Indel frequencies of TnpB reRNA and protein mutants at *PDS1*-1 and *PDS1*-2 sites in *N. benthamiana*. Data represent mean ± s.e.m. (n≥2 independent agroinfiltrations). NC indicates negative control.

**Extended Data Figure 9.**
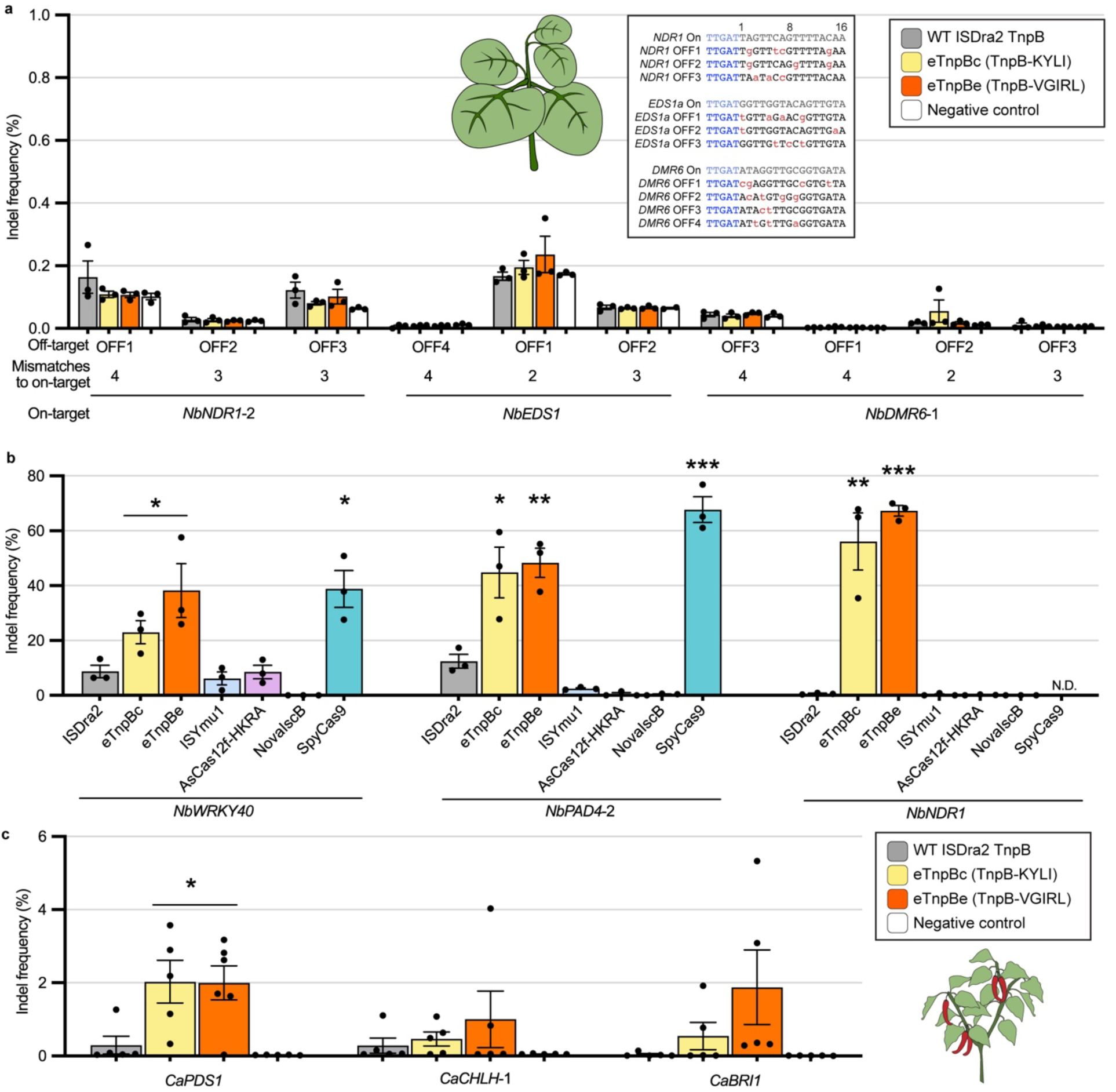
eTnpBc and eTnpbe are specific, highly active RNA-guided endonucleases in *N. benthamiana* and also show activity in pepper. **a,** Indel frequency of eTnpBc and eTnpBe compared to wild-type ISDra2 TnpB and a negative control (untransformed *Agrobacterium* infiltration) at Cas-OFFinder-predicted off-target sites in *N. benthamiana*. **b,** Indel frequencies of ISDra2 TnpB variants compared to wild-type ISYmu1TnpB, AsCas12f-HKRA, NovaIscB, and SpyCas9. **c,** Indel frequencies at three genomic sites in pepper. Data are plotted as the mean and s.e.m. from biological replicates (n≥3) in **a-c.** Stars indicate a statistically significant increase in indel frequencies compared to WT ISDra2 TnpB as calculated using a two-sided unpaired Student’s t-test. (Significance: *, **, *** for p ≤ 0.05, 0.01, 0.001, respectively) in **b-c**.

**Extended Data Figure 10.**
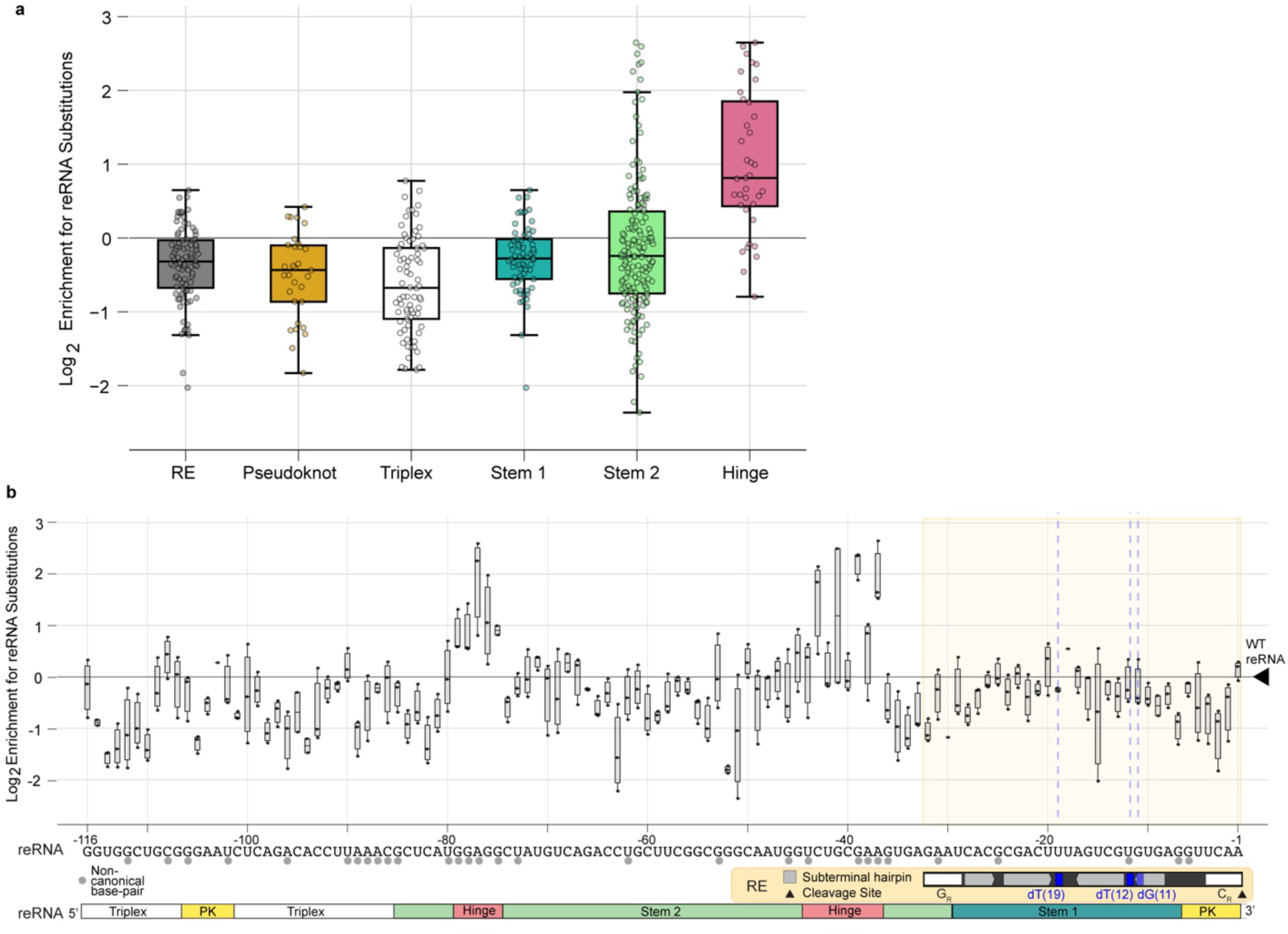
Deep mutational scanning of reRNA reveals mutational tolerance within the reRNA stem 1-RE overlap. **a,** Box and whisker plots showing the distribution of log2 enrichment for single nucleotide substitutions around the median, normalized to the wild-type reRNA log2 enrichment. Nucleotide substitutions were grouped by reRNA region. Data are represented as the median ± IQR. **b,** Log2 enrichment of single-nucleotide substitutions at each position in the reRNA, presented as a box-and-whisker plot, with each point representing an individual mutant. The x-axis includes annotations for both the overlapping RE DNA and reRNA sequences. Within the RE, key functional elements are highlighted, including the ssDNA subterminal hairpins recognized by TnpA for transposon excision, as well as the tetranucleotide cleavage (CR) and guide (GR) sequences, which form base-pair interactions and direct TnpA cleavage^42,55^. Additionally, positions within the subterminal hairpin important for TnpA binding and strand discrimination are indicated in blue^41^.

## Methods

### Deep mutational libraries construction

The TnpB reRNA DMS library was constructed from an oligonucleotide pool from Twist Bioscience, and covered the 116 nt reRNA scaffold with flanking primer-binding sites for PCR amplification. The reRNA scaffold library contained ∼600 variants, including all nucleotide substitutions; single and double nucleotide deletions; a set of double mutations in the pseudoknot; and stable tetraloop replacements in the disordered reRNA regions. The oligonucleotide library was amplified using KAPA HiFi HotStart ReadyMix with an initial denaturation at 95℃ for 3 minutes, 16 cycles of 98℃ for 20 seconds, 64℃ for 15 seconds, and 72℃ for 45 seconds, and a final extension at 72℃ for 1 minute. The amplified library was cloned into an intermediate storage vector with NEBuilder HiFi DNA Assembly Master Mix. The reRNA library was then assembled with a destination vector containing the variable 3’ reRNA sequence by Golden Gate cloning with BsaI-HFv2 and MlyI.

The reRNA plasmid library was digested with KpnI and ApaI and barcoded by assembly (NEBuilder HiFi DNA Assembly Master Mix) with ssDNA oligonucleotides with internal 15xN barcodes. The barcoded plasmid assembly was transformed into TransforMax EC100D pir-116 Electrocompetent *E. coli* and bottlenecked such that a larger culture for plasmid purification was inoculated with ∼2.4×10^4^ transformed cells (∼40 barcodes x ∼600 variants), estimated from colony-forming units (CFUs) counted from titer plates. Control plasmids containing WT and catalytically dead ISDra2 TnpB were barcoded similarly.

The ISDra2 TnpB protein sequence was codon optimized for expression in *S. cerevisiae* and mammalian cells and divided into six segments of 204 bp. For each segment, mutations for all single amino acid changes and stop codons were designed and purchased as oligonucleotide pools from Twist Bioscience with flanking primer binding sites, for a total of 8116 variants. To account for enrichment of truncations at all stop codons, the NLS tag was positioned at the N-terminus of the TnpB protein in the protein libraries, instead of the C-terminus, where it was placed in the reRNA library. Mutations were designed using the most common *S. cerevisiae* codons, except in cases where this would create a restriction site that would interfere with library cloning or plasmid linearization. In these cases, an alternative common codon set was used to introduce the intended mutation. The first methionine was excluded from mutagenesis. The six sub-libraries were amplified using KAPA HiFi HotStart ReadyMix with an initial denaturation at 95℃ for 3 minutes, 18 cycles of 98℃ for 20 seconds, 64-67℃ for 15 seconds, and 72℃ for 45 seconds, and a final extension at 72℃ for 1 minute. Amplified sub-libraries were assembled by Golden Gate cloning with BsaI-HFv2 and six corresponding intermediate cloning vectors containing the flanking WT ISDra2 sequence. Each sub-library plasmid pool was digested with BsmBI-v2, and the concentration of the digested full-length ISDra2 protein coding sequence for each sub-library was measured using a Qubit 4 Fluorometer. Each digested sub-library was mixed at an equimolar ratio and inserted into the destination vector.

Single-stranded 30xN barcode sequences were cloned into the NotI-HF- and XhoI-digested plasmid library with NEBuilder HiFi DNA Assembly Master Mix. Assemblies were transformed into TOP10 cells, and a larger culture for plasmid purification was inoculated with ∼2×10^5^ transformed cells (∼24 barcodes x ∼8100 variants).

### Combinatorial library construction

Two combinatorial libraries with an average of ∼3 and ∼5 mutations per variant were created using nicking mutagenesis, as previously described^1,2^. DNA oligos covering 19 amino acid positions and 33 possible mutations in the TnpB protein were phosphorylated and pooled in an equimolar ratio. For synthesis of the second strand, 5 pmols and 50 pmols of the phosphorylated oligo pool were initially added with 0.38 fmols of the ssDNA template plasmid, and 4.3 pmols and 43 pmols of the phosphorylated oligo pool were spiked in three times following 5 cycles of amplification, to generate the lower and higher mutation-frequency libraries, respectively.

RRYNx25RY and YYRNx25YR barcodes were cloned into the lower and higher mutation-frequency plasmid libraries, respectively, by assembly with ssDNA oligos, as described in the protein DMS library barcoding. The barcoded library assemblies were bottlenecked to ∼5×10^4^ barcoded variants with an average of ∼15 barcodes per variant. The lower and higher mutation-frequency libraries were combined in a 1:12 ratio along with barcoded, catalytically inactivated TnpB protein controls prior to transformation into yeast.

### Variant-barcode mapping

After library construction, variants were associated with their barcodes using long-read sequencing (PacBio Sequel II for the protein DMS library and Nanopore MinION for the reRNA DMS and combinatorial libraries). All reads were aligned to a reference plasmid and barcode sequences extracted using Minimap2 and SAMtools^3,4^. Sub-alignments were made for all reads with a given barcode, and a consensus sequence was created using SAMtools for all barcodes with at least two reads for PacBio sequencing and at least 10 reads for nanopore sequencing. Barcodes of incorrect length and consensus sequences containing non-programmed mutations were discarded. For the reRNA, protein, and stacked libraries, 606, 7766, and 6592 variants were mapped respectively, with an average of 33, 28, and 15 barcodes per variant respectively. Full analysis scripts and processed data available on github.

### Reporter yeast strain creation

Yeast *ade2*^-^ reporter strains were created with the *delitto perfetto* approach^5^. An intermediate *ADE2* knockout was derived from S*accharomyces cerevisiae* BY4741 (ATCC 201388, Meyen ex E.C. Hansen) using the CORE cassette GSKU, excluding Gal-I-SceI, as previously described. To create reporter strains, the intermediate strain was co-transformed with linearized DNA containing the target site flanked by duplicate homology regions, and with plasmid carrying SpyCas9 targeting the CORE cassette. SpyCas9 was constitutively expressed, triggering DSBs in the CORE cassette and repair with the linear DNA template. Target site integration was confirmed by PCR amplification and sequencing, and the strain was cured of the Cas9 plasmid. Using this approach, we generated *ade2*^-^ reporter strains UniPAM1, UniPAM2 (target 1 strain), and UniPAM5 (target 2 strain).

### Yeast pooled library selection assays

Plasmid DNA was linearized by PaqCI digestion prior to transformation, and expression vectors were assembled by gap repair homologous recombination in yeast. Linearized plasmid libraries and backbone plasmid were transformed in a 1:3 molar ratio. For each experimental replicate, 4-5 µgs of total linearized plasmid containing the DMS or combinatorial libraries of the TnpB protein was transformed. For the reRNA DMS library, 1.5 µgs total of linearized plasmid library was transformed. Yeast were transformed with lithium acetate/single-stranded carrier DNA/PEG method^6^.

After transformation, cells were resuspended in synthetic complete drop-out media lacking leucine (SCD-leucine) to select for transformation and gap repair of plasmids, and recovered overnight at 30℃. The following morning, a fraction of the culture was removed for a pre-induction time point. The remaining cells were induced in leucine liquid media with 2% galactose(w/v) at an initial OD_600_ of 1.0. Induced cultures were allowed to grow at 30℃, and culture samples were removed at multiple time points, pelleted and washed in mQ water, and plated on selective (-adenine -leucine) and non-selective (+adenine -leucine) plates. Several cell concentrations were plated on bioassay dishes (Thermo Fisher) at each time point to ensure maximal library coverage. Before plating, all cultures were grown with supplemental (160mg/ml) adenine. Yeast plates were incubated at 30℃ for 48 hours, after which colonies were scraped and plasmid DNA extracted using Zymoprep Yeast Plasmid Miniprep II. Barcodes were amplified from plasmid DNA using KAPA HiFi HotStart ReadyMix with 6-12 cycles for PCR1 and 10 cycles for PCR2. PCRs were cleaned up with Ampure XP beads (Beckman Coulter) and submitted for 150 bp paired-end sequencing on Illumina NextSeq sequencer at the IGI NGS sequencing core.

### Variant enrichment calculations

Barcode enrichment was assessed by calculating the log ratio of reads containing a given barcode in selective and nonselective samples. Barcodes with fewer than five reads in selective or non-selective conditions were removed from analysis and the log ratio was normalized by the total number of reads in selective and nonselective sequencing samples. For the protein and stacked-protein libraries, variant enrichment was calculated as the median barcode enrichment for all barcodes associated with a given variant. For the reRNA library, variant enrichment was calculated as the mean of all barcode enrichments as this produced higher replicate correlation. Variant enrichments were normalized to WT such that WT has an enrichment value of zero. A two-sided Mann-Whitney test was performed to calculate the statistical significance and effect size for each variant for each replicate.

### Yeast cleavage assays

To compare the activities of TnpB variants and orthologs, as well as of CRISPR-Cas effectors, yeast cells were transformed with 0.5 µg-1.5 µg clonal plasmids or linearized DNA for assembly of clonal plasmids, and induced as described above. At the pre-induction and post-induction timepoints, approximately 1-3 OD_600_ units were removed from the transformed yeast culture, washed with mQ water, resuspended in 200 µL mQ water, and serially diluted in triplicate. Serial dilutions were plated on selective (-adenine -leucine) and non-selective (+adenine -leucine) solid media 8 hours post-induction, unless otherwise specified. Plates were incubated at 30℃ for 48 hours, after which colony counts from serial dilutions were used to estimate the total number of colony-forming units (CFUs). Colony reversion was calculated by dividing the number of CFUs on selective media over the number of CFUs on nonselective media, multiplied by 100 for percentage.

### TnpB protein Western blots in yeast

At 24 hours post-induction of a yeast cleavage assay, 2.5 OD_600_ units of yeast cells were harvested for Western blots at 3000 x *g* for 5 minutes. Cells were resuspended in 100 μl mQ water before being lysed by adding 100 μl 0.2M NaOH and incubating at room temperature for 5 minutes. Cell lysis was pelleted by centrifugation at 21,000 x *g* for 2 minutes, washed with 200 μl of 1X PBS, and pelleted again at 21,000 x *g* for 2 minutes. Pellets were resuspended in 30 μl 1X PBS and 30 μl 4x Laemmli buffer (62.5 mM Tris-HCl, pH 6.8, 10% glycerol, 1% LDS, 0.005% Bromophenol Blue) (BioRad). Samples were boiled at 95°C for 3 minutes before 12 μl of supernatant was loaded for SDS-PAGE on a 4-20% Criterion TGX Precast Midi Protein Gel and separated at 125V for 60 minutes. Transfer to a Bio-Rad Trans-Blot Turbo Midi PVDF Transfer Pack was performed with the Trans-Turbo Turbo Transfer System. The membrane was blocked with 5% milk in TBST for 1 hour at room temperature, and then incubated overnight with mouse anti-FLAG (1:10000, Sigma F1804) and rabbit anti-PGK-1 (1:30000) in 2.5% milk in TBST overnight at 4℃. The membrane was washed three times with TBST for 10 minutes each at room temperature, then incubated with goat anti-mouse and anti-rabbit secondary antibodies (1:10000, LiCor 926-32210) for one hour at room temperature. The membrane was washed again as described before. Images were acquired on the LI-COR Odyssey CLx and processed by using ImageStudio.

### Comparative sequence and structure alignments

A multiple sequence alignment (MSA) was created using a previously described pipeline^7^. The ISDra2 TnpB protein sequence was used as the Jackhmmer query for 5 search iterations against the UniRef90 database with a domain and sequence bit score threshold of 0.1. Redundant sequences were removed using HHfilter and sequences with less than 50% coverage were removed. Per position amino acid frequencies were determined by calculating the ratio of the amino acid’s occurrence to the total number of sequences with a residue present at that position, as previously described^8^.

Structural alignments between ISDra2, ISYmu1, and ISAba30 TnpBs were generated with FoldMason^9^ (shown) and pairwise alignments^10^ with the AlphaFold2-predicted WT amino acid sequences^11^. Similarity was calculated with a Blosum62 matrix with a threshold of 1^12^.

### Mammalian genome editing

Mammalian cells (HEK393T or HEK293T-GFP) were grown in Dulbecco’s Modified Eagle Medium with high glucose, GlutaMAX supplement, and pyruvate (Thermo Fisher) supplemented with 10% fetal bovine serum (Avantor Seradigm) at 37℃ and 5% CO_2_. Cells were seeded at approximately 10,000 cells per well in 96-well plates 16-24 hours before transfection. The transfection mix was prepared by combining plasmids encoding the protein and reRNA/sgRNA (100 ng carrying TnpB, 145 ng carrying SpyCas9 for 2.6 fmols/transfection) with 9µL Opti-MEM I Reduced Serum Medium (Thermo Fisher) and 0.3µL TransIT-293 per transfection. Transfection mixes were incubated at room temperature for 30 minutes and added dropwise to the cells.

For flow cytometry, transfected plates were passaged two days post-transfection and then harvested for flow cytometry after two to five days. Cells were trypsinized with 30 µL 0.25% trypsin+EDTA was added to cells for 5 minutes at 37℃, and quenched with 120 µL 1X PBS. Cells were transferred to 96-well round-bottom plates and analyzed by flow cytometry on an Attune NxT Flow Cytometer with an autosampler. Data was analyzed using FlowJo software.

For sequencing, cells were harvested 4 days after transfection, and lysed with QuickExtract (Lucigen) according to manufacturer instructions. Lysate was used directly for polymerase chain reaction (PCR). PCR products were cleaned with Ampure XP beads (Beckman Coulter), analyzed by a 4150 TapeStation (Agilent), and submitted for 150 bp or 300 bp paired-end sequencing on MiSeq or NextSeq sequencer at the IGI NGS sequencing core. The frequencies of the mutations were assessed by CRISPResso2^13^.

### Off-target analysis

To assess the specificity of TnpB and TnpB variants, CRISPR RGEN Tools (Cas-OFFinder, http://www.rgenome.net/cas-offinder/) was used to predict potential genomic off-target sites containing the “TTGAT” TAM/PAM and 4-6 mismatches in the target sequence^14^. Primers were designed using NCBI Primer-BLAST (https://www.ncbi.nlm.nih.gov/tools/primer-blast/).

#### Nicotiana benthamiana editing

All plasmid vectors were delivered to *Nicotiana benthamiana* by *Agrobacterium tumefaciens* strain GV310 infiltration. Cultures containing the vector of interest were grown in lysogeny broth (LB) medium supplemented with kanamycin (50 µg/ml), gentamicin (100 µg/ml), and rifampicin (100 µg/ml) overnight at 30℃. The next day, cultures were spun down at 3500 x *g* for 10 minutes. The pellet was then resuspended in infiltration media (10 mM MgCl_2_, 10 mM MES (pH 5.6), 150 µM acetosyringone, in mQ H_2_O) and diluted to an OD_600_ of 1.0. The resuspension was incubated at room temperature for 3 hours before infiltration.

Syringe infiltration was performed on the lower side of 4-week old *N. benthamiana* plants on the fourth or fifth leaves from the top of the plant. Infiltrated plants were then watered and transferred back to the plant growth chamber (Percival) (16/8-h light/dark photoperiod, 80 µmol m^-2^ light intensity, 50% humidity, at 23℃) for 4 days. After the 4-day period, leaf discs were taken for each infiltrated leaf using a hole puncher tool (Electron Microscopy Sciences, 6903950). The harvested leaf tissue was lysed in 700 µL of 2% CTAB (10g CTAB, 100mM Tris HCl, 20mM EDTA, 1.4M NaCl, 1% Polyvinylpyrrolidone) or 20µL Phire Plant Direct PCR Dilution Buffer (Thermo Scientific, F160S), following flash-freezing in liquid nitrogen. Leaf lysate was used directly for PCR reactions or used for genomic DNA extraction as previously described by Yu *et al.* 2019^15^.

PCR reactions were performed using Phire Plant Direct PCR Master Mix (Thermo Fischer) and PCR products were cleaned with Ampure XP beads (Beckman Coulter), analyzed by a 4150 TapeStation (Agilent), and submitted for 150 bp or 300 bp paired-end sequencing on MiSeq or NextSeq sequencer at the IGI NGS sequencing core. The frequencies of the mutations were assessed by CRISPResso2^13^.

### Rice editing

Transgenic callus tissues and plants were generated by *Agrobacterium*-mediated transformation as previously described^16^ with minor modifications. Mature seeds of rice (*Oryza. sativa* L. *japonica* cv. Kitaake) were dehulled and surface-sterilized for 3 min with 70% (v/v) ethanol followed by 15 min in 20% (*v*/*v*) commercial bleach (5.25% sodium hypochlorite *v*/*v*) containing 1 drop of Tween 20. Seeds were washed three times with sterile water to remove residual bleach. Sterilized seeds were placed on callus induction medium (CIM)^16^ without BAP and incubated in the dark at 28°C to initiate callus induction. High-quality calli were selected and transferred to fresh CIM for proliferation.

Fifty pieces of 6- to 8-week-old calli, approximately 2-3 mm in diameter, were dried on empty sterile Petri dishes for 30 min prior to incubation with an *A. tumefaciens* AGL1 suspension (OD_600nm_ = 0.2) carrying each transformation vector. TnpB and reRNA were cloned into the pKb-TnpB2 vector previously utilized by Karmakar *et al.* to deliver wild-type ISDra2 TnpB to rice callus^17^ After a 30-min incubation, the *Agrobacterium* suspension was removed. Calli were then placed on sterile filter paper, transferred to cocultivation medium^16^ and incubated in the dark at 21°C for 3 d. Calli were then transferred to resting medium^18^ [OsCIM2 supplemented with 150 mg/L cefotaxime and 100 mg/L timentin] and incubated in the dark at 28°C for 7 d. Calli were then transferred to the selection medium (resting medium plus 40 mg/L hygromycin B) and incubated in the dark at 28°C. Tissues were transferred to the fresh selection medium every two weeks. When a sufficient amount of putatively transformed tissues was obtained after the third-round selection, a portion of healthy and well-proliferating callus tissues from each transgenic event was sampled for genomic DNA extraction.

Callus tissue (100 mg) was flash frozen in liquid nitrogen and homogenized via bead beating for 1-2 minutes at 1500 RPM. To extract genomic DNA, 350 µL 2% CTAB (10g CTAB, 100mM Tris HCl, 20mM EDTA, 1.4M NaCl, 1% Polyvinylpyrrolidone) was added to the homogenized callus and incubated at 65°C for 30 minutes. Genomic DNA was isolated with chloroform phase separation, precipitated in isopropanol, washed in 70% ethanol, dried, and resuspended in nuclease-free water.

The extracted genomic DNA was used directly for polymerase chain reaction (PCR). PCR products were cleaned with Ampure XP beads (Beckman Coulter), analyzed by a 4150 TapeStation (Agilent), and submitted for 300 bp paired-end sequencing on MiSeq or NextSeq sequencer at the IGI NGS sequencing core. The frequencies of the mutations were assessed by CRISPResso2^13^.

### Pepper editing

Serrano Tampiqueno Pepper (*Capsicum annuum* L.) was grown in a growth chamber set at 24°C with a 12-hour light and 12-hour dark cycle, a light intensity of 100 μE m-2 sec-1, and 50% humidity. The *Agrobacterium* GV3101 strain, carrying various TnpB and reRNA with target sequences, was grown in LB medium supplemented with SRG (spectinomycin at 50 μg/mL, rifampicin at 25 μg/mL, and gentamicin at 50 μg/mL) at 28°C for 14-16 hours. *Agrobacterium* cells were pelleted by centrifugation at 3500×g for 15 minutes, resuspended in infiltration medium containing 10 mM MgCl_2_, 10 mM 2-(N-morpholine)-ethanesulfonic acid (MES), and 250 μM 3′,5′-dimethoxy-4′-hydroxyacetophenone (acetosyringone) in Milli-Q water to an OD_600_ of 1.0, and incubated with gentle shaking for at least 3 hours at room temperature. *Agrobacterium* was infiltrated into two fully expanded cotyledons of 20-day-old pepper seedlings using a 1 mL needleless syringe. For CaCHLHt4 and CaBRI1t11 infiltration, *Agrobacterium* with RNA silencing suppressor p19 was added at an OD_600_ of 0.05. Infiltrated plants were kept on growth light carts, and four days later, leaf samples were collected and crushed in 35 μL of dilution buffer included in the Phire Plant Direct PCR kit (Thermo Scientific), then centrifuged at 14,000 rpm in an Eppendorf centrifuge for 5 minutes and stored at -80°C, or used directly for genotyping.

2 µL leaf extract was used for PCR with Phire Plant Direct PCR MasterMix (Thermo Scientific) according to manufacturer instructions. PCR products were cleaned with Ampure XP beads (Beckman Coulter), analyzed by a 4150 TapeStation (Agilent), and submitted for 150 bp or 300 bp paired-end sequencing on MiSeq or NextSeq sequencer at the IGI NGS sequencing core. The frequencies of the mutations were assessed by CRISPResso2^13^.

## Data availability

Sequences for MSAs were taken from UniRef90. Raw sequencing reads will be accessible on the NCBI SRA at the point of publication. Additional relevant materials (for example, plasmids and proteins) are available from the corresponding author upon reasonable request or from Addgene.

## Code availability

All code for this paper will be available at https://github.com/SavageLab/tnpb_dms.

